# Sperm Histone H3 Lysine 4 tri-methylation serves as a metabolic sensor of paternal obesity and is associated with the inheritance of metabolic dysfunction

**DOI:** 10.1101/2021.09.09.459659

**Authors:** Anne-Sophie Pepin, Christine Lafleur, Romain Lambrot, Vanessa Dumeaux, Sarah Kimmins

**Author notes:** **Summary sentence:** Paternal obesity impacts sperm H3K4me3 and is associated with placenta, embryonic and metabolic outcomes in descendants. **Declaration of interest:** none.

## Abstract

**Objective:** Parental environmental exposures can strongly influence descendant risks for adult disease. How paternal obesity changes the sperm chromatin leading to the acquisition of metabolic disease in offspring remains controversial and ill-defined. The objective of this study was to assess: (1) whether obesity induced by a high-fat diet alters sperm histone methylation; (2) whether paternal obesity can induce metabolic disturbances across generations; (3) whether there could be cumulative damage to the sperm epigenome leading to enhanced metabolic dysfunction in descendants; and (4) whether obesity-sensitive regions associate with embryonic epigenetic and transcriptomic profiles. Using a genetic mouse model of epigenetic inheritance, we investigated the role of histone H3 lysine 4 methylation (H3K4me3) in the paternal transmission of metabolic dysfunction. This transgenic mouse overexpresses the histone demethylase enzyme KDM1A in the developing germline and has an altered sperm epigenome at the level of histone H3K4 methylation. We hypothesized that challenging transgenic sires with a high-fat diet would further erode the sperm epigenome and lead to enhanced metabolic disturbances in the next generations.

**Methods:** To assess whether paternal obesity can have inter- or transgenerational impacts, and if so, to identify potential mechanisms of this non-genetic inheritance, we used wildtype C57BL/6NCrl and transgenic males with a pre-existing altered sperm epigenome. To induce obesity, sires were fed either a control or high-fat diet (10% or 60% kcal fat, respectively) for 10-12 weeks, then bred to wildtype C57BL/6NCrl female fed a regular diet. F_1_ and F_2_ descendants were characterized for metabolic phenotypes by examining the effects of paternal obesity by sex, on body weight, fat mass distribution, the liver transcriptome, intraperitoneal glucose and insulin tolerance tests. To determine whether obesity altered the F_0_ sperm chromatin, native chromatin immunoprecipitation-sequencing targeting H3K4me3 was performed. To gain insight into mechanisms of paternal transmission, we compared our sperm H3K4me3 profiles with embryonic and placental chromatin states, histone modification and gene expression profiles.

**Results:** Obesity-induced alterations in H3K4me3 occurred at genes implicated in metabolic, inflammatory, and developmental processes. These processes were associated with offspring metabolic dysfunction and corresponded to genes enriched for H3K4me3 in embryos, and overlapped embryonic and placenta gene expression profiles. Transgenerational susceptibility to metabolic disease was only observed when obese F_0_ had a pre-existing modified sperm epigenome. This coincided with increased H3K4me3 alterations in sperm and more severe phenotypes affecting their offspring.

**Conclusions:** Our data suggest sperm H3K4me3 might serve as a metabolic sensor that connects paternal diet with offspring phenotypes via the placenta. This non-DNA based knowledge of inheritance has the potential to improve our understanding of how environment shapes heritability and may lead to novel routes for the prevention of disease. This study highlights the need to further study the connection between the sperm epigenome, placental development and children’s health.

## 1 Introduction

The prevalence of obesity and type II diabetes is growing globally at rates indicating that environment rather than genes is the principal driver. Exposures to high-fat diet, toxicants or micronutrient deficiency can impact our health and that of future generations [1–4]. Only now are we beginning to identify mechanisms linking these exposures to parental and offspring health. One connection between environment and health is the epigenome. The epigenome refers to the biochemical content associated with DNA that impacts gene expression and chromatin organization. Uncovering how genomic information is organized and regulated through epigenetic processes to control gene expression and cell functions in the next generation is still in a nascent stage. We and others have shown that errors in epigenomic profiles in sperm can be induced by environmental exposure to toxicants such as those in insecticides and plastics, obesity, and poor diet [5–10]. We recently demonstrated that these epigenome changes at the level of chromatin can be transmitted via sperm to alter embryonic gene expression, development, and offspring health [9]. Historically, parental health and fertility have focused predominantly on the mother, although it is clear a father’s health and lifestyle can also impact his children’s health. How epimutations in sperm functionally impact the embryo urgently require elucidation to prevent transmission of disease from father to offspring.

Metabolic disease including obesity and type II diabetes can in part be attributed to genetic factors with a 5-10% increased risk [11]. The remaining risk is attributable to environmental-epigenetic interactions including potentially those of our ancestors. This possibility is supported by epidemiological and animal studies. Transgenerational effects are suggested by studies in humans that linked the food supply of grandfathers to obesity and cardiovascular disease in their grandchildren [12–14]. However, the ability for diet to induce transgenerational effects in animal models remains controversial and requires more in depth studies addressing the underlying molecular mechanisms [15–17]. To date, studies using mice to assess the impact of diet and obesity in relation to the sperm epigenome, have focused on the DNA methylome and noncoding RNA (ncRNA) as the potential sperm-borne mediators of metabolic disease [6,18–23]. The role of sperm chromatin in the non-genetic inheritance of metabolic disorders is unknown. In human and mouse sperm, histone H3 lysine 4 trimethylation (H3K4me3) localizes to genes involved in metabolism and development [9,24,25]. Moreover, sperm H3K4me3 can be altered by folate deficiency and influences embryonic development and gene expression [15, 26]. This association of histone modifications in sperm with offspring phenotypes has since been confirmed in other mouse models [27, 28]. Based on these observations, we hypothesized that sperm H3K4me3 may serve as a metabolic sensor that is implicated in the paternal transmission of obesity-associated disease in offspring.

A focus of this study was to identify whether paternal obesity impacts the F_1-_F_2_, and if so, to identify potential mechanisms of this non-genetic inheritance. In our transgenic (TG) mouse model of epigenetic inheritance, male mice overexpress the histone demethylase KDM1A specifically in spermatogenesis, resulting in sperm with alterations in H3K4me2 and me3. Of note, only H3K4me3 has been implicated in transgenerational inheritance in this mouse model [26]. Therefore, as sperm H3K4me3 is responsive to paternal folate deficiency [9], and has been implicated in transgenerational inheritance [26], we targeted this mark in sperm to probe in response to paternal obesity and as a potential mediator of inheritance. In this study, we aimed to: 1) assess the impact of high-fat diet (HFD) induced paternal obesity on sperm H3K4me3 and its association with metabolic dysfunction across generations, and 2) determine if descendants of obese TG sires with a previously altered sperm epigenome would show more severe metabolic dysfunction. To address these aims, we used wildtype (WT), or the germline specific KDM1A-overexpressing TG mice, in combination with a diet-induced obesity model. These TG sires have descended from males that have an altered sperm epigenome and whose ancestors had compromised health (see Materials and Methods for details). This TG model is used to represent an at-risk population that may be more susceptible to poor health when challenged with obesity. Here, we demonstrate that a paternal high-fat diet induces F_0_ obesity and metabolic dysfunction in the F_1_. Remarkably, transgenerational phenotypes were only observed in descendants of obese KDM1A TG males, and this was associated with enhanced alterations in H3K4me3 enrichment in obese TG sperm. This suggests that the risk of transgenerational disease transmission may be greater if an ancestor has had prior exposures that cause pre-existing damage to the sperm epigenome. Concordant with the metabolic phenotypes observed in offspring, obesity-induced alterations in sperm H3K4me3 occurred at genes involved in development, placenta formation, inflammatory processes, glucose and lipid metabolic pathways. These sperm altered H3K4me3 regions persist in the embryo and placenta, supporting a role for sperm H3K4me3 in paternal origins of adult-onset metabolic disorders.

## 2 Materials and Methods

### 2.1 RESOURCE AVAILABILITY

#### 2.1.1 Lead contact

Further information and requests for resources and reagents should be directed to and will be fulfilled by the Lead Contact, Sarah Kimmins.

#### 2.1.2 Materials availability

This study did not generate new unique reagents.

#### 2.1.3 Data and code availability

The sperm H3K4me3 ChIP-Seq and liver RNA-Seq data generated in this study are available at the following GEO accession number: GSE178096.

### 2.2 EXPERIMENTAL MODEL AND SUBJECT DETAILS

#### 2.2.1 Animals

All animal procedures were carried out in accordance with the guidelines of the Faculty Animal Care Committee of McGill University, Montreal. For the wildtype line (WT), C57BL/6NCrl 8-week old males and 6-week old females were purchased from Charles Rivers Laboratory and were allowed one week of acclimation before breeding. For the KDM1A transgenic line (TG), mice were generated as previously described [15], with the same genetic background as the wildtype line. The F_0_ TG mice used in this study were from the 11th generation. The earlier generations of mice in this KDM1A TG had severe developmental abnormalities, pre-implantation loss and early post-natal death [15]. Over time we have selected against the severe phenotype by breeding the mice that survive and are normal. Single males were housed with two females to generate the F_0_ generation. All animals were given access to water and food *ad libitum* and were maintained on a controlled light/dark cycle.

### 2.3 METHODS DETAILS

#### 2.3.1 Diet experiments and animal breeding

The low-fat control diet (CON; D12450J) and high-fat diet (HFD; D12492) were obtained from Research Diets, and selected based on the matched amounts of sucrose, vitamin mix and folate. Diets’ macronutrients composition are listed in Table S1. Males of the F_0_ generation were generated from at least 7 different sires per group. F_0_ males were weaned at 3 weeks of age and randomly assigned to either a CON or HFD. The number of animals per group, per sex and per generation, used for all metabolic characterization tests can be found in Table S2. Total body weights were monitored weekly. Cumulative caloric intake was recorded weekly by weighting pellets from the food hopper and calculated as kilocalorie per animal. The diet intervention spanned 10-12 weeks followed by 2 weeks of metabolic testing (at 4 months of age), 1 week of rest and 1-2 weeks of breeding with 7-week old C57BL/6NCrl females. Females used for breeding were housed with males overnight (1-2 females per male) and removed the following morning. This was repeated until a vaginal plug was detected, 3 nights per week for a maximum of 2 consecutive weeks. A limitation worth noting is that despite these precautions the females were exposed for a maximum of 6 nights to the HFD pre-pregnancy during this breeding period. However the impacts of this exposure are minimal as female mice require several weeks (∼5-8 weeks) before significant weight gain on a HFD [29].

Litter sizes (number of pups per litter) were recorded, and sex ratios (ratio of male pups over total number of pups) were calculated for all litters generated and can be found in Tables S3 and S4, respectively. The same timeline was used to generate the F_1_ and F_2_ animals. All females used for breeding and all F_1_ and F_2_ were fed a regular chow diet (2020X Teklad rodent diet, Envigo). All animals were sacrificed at 22 weeks (±2 weeks) by carbon dioxide asphyxiation under isoflurane anesthesia.

#### 2.3.2 Metabolic testing

Assessment of metabolic parameters was conducted at 4 months of age within 2 consecutive weeks according to the standard operating procedures of the National Institutes of Health (NIH) Mouse Metabolic Phenotyping Center [30]. For the glucose tolerance test, animals were fasted overnight for 15 hours (± 1 hour) starting at 6:00PM with free access to water. Blood glucose was measured before and 15, 30, 60 and 120 minutes following an intraperitoneal injection of 2 g/kg of a 20% glucose solution (D-glucose, G7021, Sigma Aldrich) with one drop of blood from the tail-tip using a glucometer (Accu-Chek Aviva Nano). For the insulin tolerance test, animals were fasted for 6 hours (± 1 hour), starting at 9:00AM with free access to water. Blood glucose was measured before and 15, 30, 60 and 120 minutes following an intraperitoneal injection of 1 IU/kg insulin (Insulin solution, I9278, Sigma Aldrich), with one drop of blood from the tail-tip using a glucometer (Accu-Chek Aviva Nano). The area under the curves (AUCs) for the tolerance tests were calculated using the trapezoidal rule (GraphPad Prism, version 8). For the baseline blood glucose levels, blood glucose levels were measured after an overnight fasting of 15 hours (± 1 hour) with one drop of blood from the tail-tip using a glucometer (Accu-Chek Aviva Nano).

#### 2.3.3 Tissue collection

At necropsy, mice were dissected to collect adipose tissue (gonadal and mesenteric white adipose depots; gWAT and mWAT, respectively) and a liver lobe (left lateral lobe or *lobus hepatis sinister lateralis* for RNA-sequencing). All tissues were weighed, transferred to a clean tube, snap frozen in liquid nitrogen and stored at -80°C until subsequent downstream experiments. Cauda epididymides were weighed and immediately used for sperm isolation.

#### 2.3.4 Sperm isolation

Spermatozoa were isolated from paired caudal epididymides [31, 32]. Cauda epididymides were cut into 5 mL of freshly-prepared Donners medium (25 mM NaHCO_3_, 20 mg ml^-1^ BSA, 1 mM sodium pyruvate, 0.53% vol/vol sodium DL-lactate in Donners stock) and gently agitated to allow to swim out for 1 hour at 37°C. The solution was passed through a 40-µm cell strainer (Fisher Scientific, #22363547) and washed three times with phosphate-buffered saline (PBS). The swim out and the cleaning steps remove 99% of contaminating somatic cells which is visually confirmed and has been validated in our prior studies [9,15,24,26,31,32]. The sperm pellet was cryopreserved in freezing medium (Irvine Scientific, cat. #90128) and kept in a -80°C freezer until the chromatin immunoprecipitation experiment.

#### 2.3.5 RNA-Sequencing and library preparation

RNA extraction was performed using the RNeasy Mini Kit (Qiagen, cat. #74104) following the manufacturer’s protocol with slight modifications. In brief, 15-20 mg of liver lobes were cut on dry ice using a sterile scalpel and Petri dish. Samples were lysed in 350 µL of a denaturing buffer (*Buffer RLT* with beta-mercaptoethanol) and homogenized with homogenizer pestles. Lysates were centrifuged at maximum speed for 3 minutes and the supernatants transferred to a clear tube. Ethanol (50%) was added to lysates to promote selective binding of RNA molecules to the silica-based membrane when applied to the spin columns. To avoid genomic DNA contamination, an additional DNase digestion was performed. Finally, membranes of the spin columns were washed twice with 500 µL of *Buffer RPE* and total RNA was eluted using 30 µL of RNase-free water. Libraries were prepared and sequenced at the *Génome Québec Innovation Centre* with single-end 50 base-pair (bp) reads on the illumina HiSeq 4000 and paired-end 100 bp reads on the illumina NovaSeq 6000 S2 sequencing platforms.

#### 2.3.6 ChIP-Sequencing and library preparation

Chromatin immunoprecipitation was performed as we have previously described [31, 32]. In brief, spermatozoa samples in freezing media were thawed on ice and washed with 1 mL phosphate-buffered saline. For each sample, two aliquots of 10 µL were used to count spermatozoa in a hemocytometer under microscope, and 10 million spermatozoa were used per sample (n=5 sample per group). Sperm chromatin was decondensed in 1 M dithiothreitol (DTT; Bio Shop, #3483-12-3) and the reaction quenched with N-ethylmaleimide (NEM). Samples were lysed in lysis buffer (0.3 M sucrose, 60 mM KCl, 15 mM Tris-HCl pH 7.5, 0.5 mM DTT, 5 mM McGl_2_, 0.1 mM EGTA, 1% deoxycholate and 0.5% NP40). An MNase enzyme (15 units; Roche, #10107921001) was added to aliquots containing 2 million spermatozoa in an MNase buffer (0.3 M sucrose, 85 mM Tris-HCl pH 7.5, 3 mM MgCl_2_ and 2 mM CaCl_2_), for exactly 5 minutes at 37°C. The digestion was stopped with 5 mM EDTA. Samples were centrifuged at maximum speed for 10 minutes, and the supernatants of aliquots from each sample were pooled back together. Each tube was supplemented with a protease inhibitor to obtain an 1X solution (complete Tablets EASYpack, Roche, #04693116001). Magnetic beads (DynaBeads, Protein A, Thermo Fisher Scientific, #10002D) were pre-blocked in a 0.5% Bovine Serum Albumin (BSA, Sigma Aldrich, #BP1600-100) solution for 4 hours at 4°C and then used to pre-clear the chromatin for 1 hour at 4°C. Pulling down of the pre-cleared chromatin was performed with the use of magnetic beads that were previously incubated with 5 µg of antibody (Histone H3 Lysine 4 trimethylation; H3K4me3; Cell Signaling Technology, cat. #9751) for 8 hours at 4°C. Immunoprecipitation of the chromatin with the beads-antibody suspension was performed overnight at 4°C. Beads bound to the chromatin were subjected to a 3-step wash, one wash with Washing Buffer A (50 mM Tris-HCl pH 7.5, 10 mM EDTA, 75 mM NaCl) and two washes with Washing Buffer B (50 mM Tris-HCl pH 7.5, 10 mM EDTA, 125 mM NaCl). The chromatin was eluted in 250 µL of Elution Buffer (0.1 M NaHCO_3_, 0.2% SDS, 5 mM DTT) by incubating the beads twice (2 x 125 µL) shaking at 400 rpm for 10 minutes at 65°C, vortexing vigorously and transferring the chromatin elute in a clean tube. The eluted chromatin was finally treated with 5 µL of RNase A (Sigma Aldrich, #10109169001) by shaking in a thermomixer at 400 rpm for 1 hour at 37°C, and then with 5 µL of Proteinase K (Sigma Aldrich, #P2308) overnight at 55°C. DNA was extracted and purified using the *ChIP DNA Clean and Concentrator kit* (Zymo Research, #D5201) using the manufacturer’s protocol, eluted with 25 µL of the provided elution buffer. Size selection of the mononucleosomes (147 bp) was performed with the use of Agencourt AMPure XP beads (Beckman Coulter, #A63880). Libraries were prepared in-house using the Ultra-low Input Library kit (Qiagen; #180495). Libraries were sequenced with single-end 50 bp reads on the illumina HiSeq 4000 sequencing platform (n=5 samples per experimental group).

#### 2.3.7 Pre-processing

##### 2.3.7.1 Liver RNA-Sequencing data

All samples were processed with the same parameters with the exception of those sequenced on the NovaSeq platform to adapt for paired-end sequencing and sequencing read length. Reads were trimmed using *Trim Galore* (version 0.5.0, parameters for HiSeq: --phred33 --length 36 -q 5 --stringency 1 -e 0.1; parameters for NovaSeq: --paired --retain_unpaired --phred33 --length 36 -q 5 --stringency 1 -e 0.1) [33]. Trimmed reads were aligned to the *Ensembl* Genome Reference Consortium mouse reference 38 (GRCm38) primary assembly using *hisat2* (version 2.1.0, parameters: -p 8 --dta) [34]. Aligned files with SAM format were converted to binary SAM format (BAM) and sorted by genomic position using *SAMtools* (version 1.9) [35]. Transcripts were assembled and gene abundances calculated using *Stringtie* (version 2.1.2, parameters: -p 8 -e -B -A) [36].

##### 2.3.7.2 Sperm ChIP-Sequencing data

Sequencing reads were trimmed using *Trimmomatic* on single-end mode to remove adapters and filter out low-quality reads (version 0.36, parameters: 2:30:15 LEADING:30 TRAILING:30) [37]. Trimmed reads were aligned to the *Mus Musculus* mm10 genome assembly using *Bowtie2* (version 2.3.4) [38]. Unmapped reads were removed using *SAMtools* (version 1.9) [35], and those with 3 mismatches or more were filtered out using *Perlcode*. BAM coverage files (BigWig) were generated using *deeptools2 bamCoverage* function (version 3.2.1, parameters: -of bigwig -bs 25 -p 20 --normalizeUsing RPKM -e 160 --ignoreForNormalization chrX) [39].

##### 2.3.7.3 Other publicly available ATAC-Sequencing or ChIP-Sequencing datasets

Raw files were downloaded from the National Centre for Biotechnology Information (NCBI) using the Sequencing Read Archive (SRA) Toolkit for 2-cell H3K4me3 ChIP-Seq [40] (GEO: GSE73952), MII oocyte H3K4me3 ChIP-Seq [41] (GEO: GSE71434), sperm ATAC-Seq [42] (GEO: GSE79230), 4-cell and morula ATAC-Seq [43] (NCBI SRA: SRP163205), TE H3K4me3 ChIP-Seq [40] (GEO: GSE73952), and placenta H3K4me3 ChIP-Seq [44] (GEO: GSE29184). Files were pre-processed as described above for the sperm H3K4me3 ChIP-Sequencing with slight modifications to adapt for datasets with paired-end reads and for different sequencing read lengths.

##### 2.3.7.4 Other publicly available RNA-Sequencing data

Raw files for 4-cell and morula [43] (NCBI SRA: SRP163205), TE [40] (GEO: GSE73952), and placenta [45] (NCBI SRA: SRP137723) RNA-Seq were downloaded from NCBI using the SRA Toolkit. Files were pre-processed as described above for the liver RNA-Sequencing with slight modifications to adapt for datasets with paired-end reads and for different sequencing read lengths.

##### 2.3.7.5 Paternal allele 2-cell embryo ChIP-Sequencing data

Raw files for 2-cell H3K4me3 ChIP-Seq [41] (GEO: GSE71434) were downloaded from NCBI using the SRA Toolkit. These datasets from mouse 2-cell embryos were generated by crossing males and females of different strains, permitting the assignment of reads to the paternal-specific allele. *SNPsplit* (version 0.3.2) was used to build a reference genome with PWK_PhJ single nucleotide polymorphism (SNPs) masked [46]. Reads were aligned to the generated PWK_PhJ SNPs N-masked reference genome using *Bowtie2* (parameters: -p 10 -t -q -N 1 -L 25 -X 2000 --no-mixed --no-discordant). Aligned files with SAM format were converted to binary SAM format (BAM) and sorted by genomic position using *SAMtools (*version 1.9) [35]. *SNPsplit* (version 0.3.2) was used to assign reads to either the paternal (PWK_PhJ) or the maternal (C57BL/6) genome based on SNPs origin. BAM coverage files (BigWig) were generated using *deeptools2 bamCoverage* function (parameters: -of bigwig -bs 25 -p 20 --normalizeUsing RPKM -e 160 --ignoreForNormalization chrX).

### 2.4 QUANTIFICATION AND STATISTICAL ANALYSIS

#### 2.4.1 Visualization and statistical analyses for metabolic characterization

Visualization of the metabolic characterization data was performed using Jupyter Notebook (version 6.0.1) with Python (version 3.7.4), with the use of the following packages: *seaborn* (version 0.9.0) [47], *numpy* (version 1.17.2) [48], and *panda* (version 0.25.2) [49]. The *pyplot* and *patches* modules were loaded from the *matplotlib* library (version 3.4.2) [50]. Statistical analyses were conducted using GraphPad Prism 8. For all tests, a p-value less than 0.05 was considered significant. To assess main effects of time, diet or genotype, and diet-genotype interactions, for the blood glucose curves of the glucose and insulin tolerance tests, and for cumulative energy intake and growth trajectories during the diet intervention, 3-way ANOVA with Geisser-Greenhouse correction was used. Significance for individual time points was tested using multiple t-test with a Holm-Sidak correction. For total body weight, mesenteric and gonadal white adipose tissue weight, baseline blood glucose and the area under the curve for the glucose and insulin tolerance tests, main effects of diet, genotype, and diet-genotype interactions were assessed using 2-way ANOVA. To assess significance for pairwise comparisons of interest, normality was assessed by D’Agostino and Pearson’s test to determine whether parametric or nonparametric statistics should be conducted. For parametric tests, an F-test was used to determine whether equal variance can be assumed. The unpaired t-test or the Welch’s t-test was used accordingly. For nonparametric tests, the Mann-Whitney test was used. Litter sizes and sex ratios were analyzed by 2-way ANOVA to assess main effects of genotype and diet, and interaction, followed by Tukey’s multiple comparisons test.

#### 2.4.2 Bioinformatics analysis

All bioinformatics analyses were conducted using R version 4.0.2 (R Core Team, 2018).

#### 2.4.3 Liver RNA-Sequencing data

Transcripts with a mean count below 10 were filtered out, conferring a total of 27,907 and 45,992 detected expressed transcripts in samples sequenced on the illumina HiSeq and NovaSeq platforms, respectively. The samples tended to cluster by RNA Integrity Number (RIN), which was corrected for in the differential analysis (Fig. S3B). Differential expression analysis was conducted using *DESeq2* (version 1.28.1) [52], by including sample’s RIN value and group in the design formula. Independent hypothesis weighting (IHW, version 1.16.0) was used to correct for multiple testing and prioritization of hypothesis testing based on covariate (i.e. the means of normalized counts) [53]. IHW calculates weight for each individual p-value and then applies the Benjamini-Hochberg (BH) procedure to adjust weighted p-values [54]. Finally, we used the Lancaster method to perform a gene-level analysis at single transcript resolution (*aggregation* package, version 1.0.1) [55]. Lancaster applies aggregation of individual transcripts p-values to obtain differentially expressed genes while capturing changes at the transcript level. Genes with a Lancaster p-value below 0.05 were considered significant.

For data visualization, transcript counts were normalized using variance stabilizing transformation without the use of blind dispersion estimation (i.e. with parameter *blind=FALSE*) [52]. This transformation approach translates data on a log_2_ scale, allows correction for library size and removes the dependence of the variance on the mean (heteroscedasticity). Variance-stabilized transcript counts were corrected for RIN values using *limma’s removeBatchEffect* function (version 3.44.3) [56]. Pearson correlation heatmaps were generated using the *corrplot* package (version 0.88) [57], with samples ordered by hierarchical clustering. Principal component analysis was performed using *DEseq’s plotPCA* function, with RIN values and sexes labeled. Heatmaps of differentially expressed genes were generated with the *Pheatmap* package (version 1.0.12) [58], with transcripts ordered by k-means clustering (n kmeans=2) and samples ordered by hierarchical clustering using complete-linkage clustering based on Euclidean distance. Alluvial plots were generated with *ggplot2* (version 3.3.3) [59], and overlap of differentially expressed genes across genotypes, generations and sexes were determined by the *GeneOverlap* package (version 1.24.0) [60], which uses a Fisher’s exact test to compute p-values.

#### 2.4.4 Visualization, Semantic similarity, and Enrichment Analysis of Gene Ontology (ViSEAGO)

Gene ontology (GO) analysis was performed using the *ViSEAGO* package (version 1.2.0) [61]. Gene symbols and *EntrezGene* IDs from the *org.Mm.eg.db* database were retrieved using the *AnnotationDbi* package. GO annotations were retrieved from *EntrezGene* for the *Mus Musculus* species (ID=”10090”) using the *ViSEAGO EntrezGene2GO* followed by *annotate* functions. ViSEAGO uses topGO to perform GO terms enrichment tests on the sets of genes of interest (differentially expressed genes). We used the Biological Process (BP) ontology category with Fisher’s exact test (classic algorithm), and a p-value below 0.01 was considered significant. Results of enrichment tests for each set of genes of interest were then merged and hierarchical clustering was performed based on Wang’s semantic similarity distance and *ward.D2* aggregation criterion. Results are visualized on a heatmap where GO terms are ordered by hierarchical clustering based on their functional similarity and GO terms enrichment significance is shown as a color gradient (-log_10_ p-value) in each set of differentially expressed genes of interest.

#### 2.4.5 Sperm ChIP-Sequencing data

To detect genomic regions enriched with H3K4me3 in sperm, we used *csaw* (version 1.22.1) [62] to scan the genome into windows of 150 bp. Windows with a fold-change enrichment of 4 over bins of 2,000 bp (background) were considered enriched. Enriched regions less than 100 bp apart were merged for a maximum width of 5,000 bp, conferring a total of 30,745 merged enriched regions. Counts in enriched regions were normalized using TMM normalization followed by *ComBat’s* correction for batch effects (*sva* package, version 3.36.0) [63, 64]. Spearman correlation heatmaps and MA-plots were generated using raw and normalized counts at enriched regions using *corrplot* (version 0.88) [57], and *graphics* packages, respectively.

Principal component analysis was conducted on normalized counts in enriched regions, by comparing WT HFD vs WT CON (effect of diet in WT), TG HFD vs TG CON (effect of diet in TG), and WT CON vs TG HFD (combined effects of genotype and HFD). Based on visual assessment of the separation of samples according to dietary or genotype groups along Principal Component 1 (PC1; x axis) or 2 (PC2; y axis), the top 5% regions contributing the PC of interest were selected. Permutational multivariate analysis of variance (PERMANOVA) was conducted to determine whether variation is attributed to dietary/genotype group, using the *adonis* function (*vegan* package, version 2.5-7) [65]. Euclidean distances were used as a metric, 999 permutations were performed, and a p<0.05 was considered significant. The directionality change in enrichment was identified based on the positive (up-regulated regions) and negative (down-regulated regions) log_2_ fold change values of the median of normalized counts using *gtools’ foldchange2logratio* function. Regions with increased and decreased enrichment for each comparison of interest were visualized using *Pheatmap* (version 1.0.12) [58]. Regions distance relative to transcription start site (TSS) were annotated and visualized using the package *chipenrich* (version 2.12.0) [66]. Gene ontology analysis was performed using *topGO* (version 2.40.0) for genes with increased or decreased H3K4me3 enrichment at the promoter region for each comparison of interest. We used the Biological Process (BP) ontology category with Fisher’s exact test *weight01Fisher* algorithm [67], and a p-value less than 0.05 was considered significant. Genomic regions with deH3K4me3 were annotated using *annotatr* (version 1.14.0) [68] including CpG annotations and basic genes genomic features. Upset plots were generated using *UpsetR* (version 1.4.0) [69], by ordering each set by frequency and displaying 12 sets. Z-scores were calculated using *regioneR’s overlapPermTest* (version 1.20.1) which performs a permutation test (n=1,000 permutations) to assess whether a set of regions is significantly enriched to a specific genomic feature compared to genomic regions from the whole genome [70]. Genome browser snapshots were generated using *trackplot* [71].

To assess linear trends associated with the cumulative exposure of KDM1A overexpression and high-fat feeding in sperm, we ran *DESeq2* (version 1.28.1) on the top 5% regions contributing to Principal Component 2 (PC2; n=1,538 regions) associated with sample separation when comparing WT CON and TG HFD normalized counts. In the design formula, we included sample’s batch information, and assigned a numerical value for each sample based on their group category (WT CON=1, WT HFD=2, TG CON=2, TG HFD=3). Independent hypothesis weighting (IHW) was used to correct for multiple testing and prioritization of hypothesis testing based on covariate (i.e. the means of normalized counts) [53]. Median of normalized counts were used to depict the increased and decreased trend of significant regions (adjusted p-value less than 0.2) across groups recoded on a numerical scale as defined above.

## 3 Results

### 3.1 Paternal obesity induces metabolic phenotypes in a sex-specific manner that are enhanced in KDM1A F_1_ and F_2_ transgenic descendants

#### 3.1.1 Impact of paternal obesity on offspring bodyweight and fat accruement

Beginning at weaning until 20 weeks, inbred C57BL/6NCrl control mice (WT), or KDM1A heterozygous transgenics (TG) were fed either a calorie-dense high-fat diet (HFD; 60% kcal fat), or a sucrose- and vitamin-matched control diet (CON; 10% kcal fat) (Fig. 1A-C and Table S1). Table S2 provides the animal numbers by sex, generation, and genotype for metabolic characterization. In the 2-4 weeks post-weaning, F_0_ males on the HFD consumed more calories and gained significantly more weight than CON males irrespective of genotype (Fig. S1A-B). These effects persisted throughout the diet intervention (Fig. S1A-C), with TG HFD males weighing the most at 4 months (Fig. S1C_i_). This trend continued in the TG male F_1_ and F_2_ descendants (fed regular chow), with weights being significantly more than the F_1_ and F_2_ of TG CON and WT HFD (Fig. S1C_ii-iii_). Indicating sex-specific responses to paternal obesity, in female descendants the changes in body weight and fat deposition differed from males (Fig. S1C-E). To assess fat accruement, we measured visceral mesenteric and gonadal white adipose tissue (mWAT and gWAT, respectively). All male (F_0_) on the HFD accumulated more mWAT compared to CON males, with no genotype effect (Fig. S1D_i_). Male and female F_1_ offspring sired by WT HFD or TG HFD had increased mWAT fat mass compared to WT CON and TG CON (Fig. S1D_ii_ and S1D_iv_, respectively). Strikingly, mWAT stores were greater in TG HFD F_1_ and F_2_ males and females compared to WT HFD descendants (Fig. S1D_ii-v_). Gonadal fat depots in F_0_ males were not impacted by the HFD (gWAT; Fig. S1E_i_), while male WT HFD F_1_ showed increased gWAT, and TG HFD F_1_ did not (gWAT; Fig. S1E_ii_). Like for body weight and mWAT, male and female F_2_ TG HFD had increased gWAT in comparison to WT HFD (Fig. S1E_v_). Overall analysis of body weight and fat accruement revealed sex-specific responses in descendants with transgenerational effects of paternal obesity being detected only in the TG HFD descendants of both males and females.

**Figure 1:**
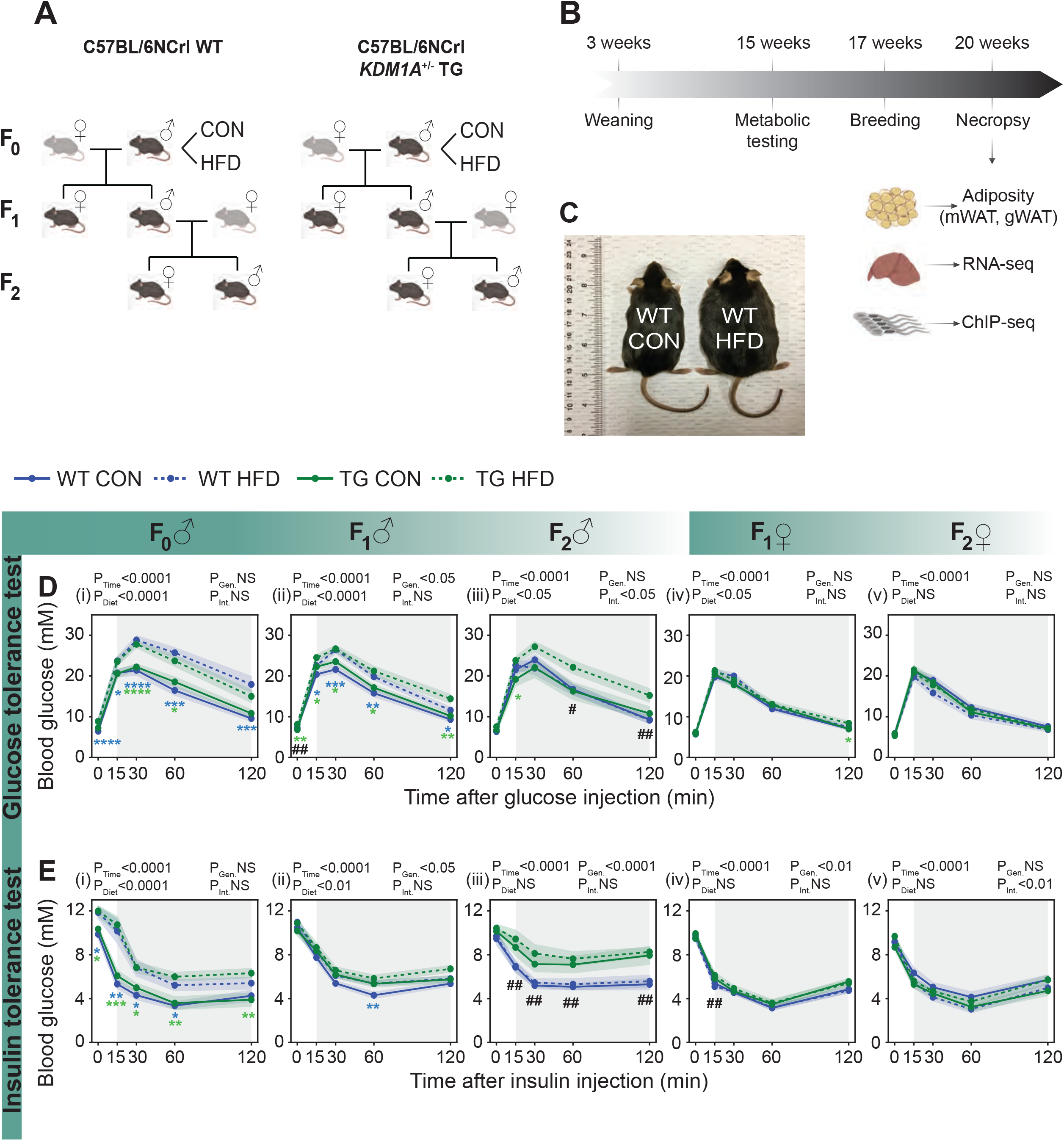
Paternal obesity induces transgenerational metabolic phenotypes in a sex-specific manner that are enhanced in KDM1A descendants. A) Experimental mouse model depicting breeding scheme and generations studied. Male C57BL/6NCrl (WT) and KDM1A^+/-^ transgenics (TG, C57BL/6NCrl) were fed either a control diet (CON) or high-fat diet (HFD) from weaning for 10-12 weeks, then mated to 8-week-old C57BL/6NCrl females fed a regular chow diet (CD). Animals studied per experimental group: F_0_ (n= 15-25 males), F_1_ (n= 28-49 per sex) and F_2_ (n= 8-21 per sex). Created with BioRender.com. B) Experimental timeline for metabolic testing and downstream experiments performed for each generation (F_0-2_). Metabolic profiles were measured after the diet intervention at 15 weeks of age and included: baseline blood glucose, and intraperitoneal glucose and insulin tolerance tests (ipGTT and ipITT, respectively). Visceral adipose depots were weighed (mWAT: mesenteric white adipose tissue and gWAT: gonadal white adipose tissue) and the left lateral lobe of the liver used for RNA-sequencing (RNA-seq). Sperm from cauda epididymides were used for chromatin immunoprecipitation followed by sequencing (ChIP-seq), targeting histone H3 lysine 4 tri-methylation (H3K4me3). Created with BioRender.com. C) Age-matched male mice fed either a control (left) or a high-fat diet (right) for 12 weeks. D) Glucose tolerance test. Blood glucose levels before and after (shaded in grey) an intraperitoneal glucose injection, after overnight fasting (15 ±1 hour) at 4 months of age in F_0_ males (i), F_1_ males (ii), F_2_ males (iii), F_1_ females (iv) and F_2_ females (v). E) Insulin tolerance test. Blood glucose levels before and after (shaded in grey) an intraperitoneal insulin injection, after a 6-hour (±1 hour) fasting at 4 months of age in F_0_ males (i), F_1_ males (ii), F_2_ males (iii), F_1_ females (iv) and F_2_ females (v). Results are shown as mean ± SEM. Significance for main effects of diet, genotype, time, and for diet-genotype interactions are shown above each graph. Significance for pairwise comparisons are shown as the following: *P<0.05, **P<0.01, ***P<0.001, ****P<0.0001 (in blue; WT CON vs WT HFD, in green; TG CON vs TG HFD) and ^#^P<0.05, _##P<0.01_ (WT HFD vs TG HFD).

#### 3.1.2 Impact of paternal obesity on glucose homeostasis

Next, we assessed glucose metabolism and insulin sensitivity by glucose tolerance (GTT), and insulin tolerance tests (ITT). These were conducted following the standard operating procedures of the NIH Mouse Metabolic Phenotyping Center [30]. First, we assessed the effects of the HFD on fasting blood glucose. Consumption of a HFD resulted in elevated baseline glucose in male (F_0_) WT HFD and TG HFD in comparison to WT CON and TG CON, respectively (Fig. S2Ai). Male TG HFD descendants (F_1_), but not WT HFD descendants had significantly elevated fasting blood glucose (Fig. S2A_ii_). In contrast, the glycemic status of all descendant females (F_1_ and F_2_) did not differ between groups (Fig. S2A_iv_-_v_). The same animals used to assess baseline glucose were then given an intraperitoneal glucose challenge and the rate of glucose disposal measured. Analysis of GTT data showed that F_0_ WT HFD and TG HFD were glucose intolerant following glucose injection in comparison to F_0_ CON males (Fig. 1D_i_). Indicating that there were intergenerational effects of paternal obesity, elevated glucose levels persisted across the GTT time-course for the F_1_ WT and TG HFD males (Fig. 1D_ii_). Interestingly, glycemic response impairments persisted in the F_2_ generation of male descendants of TG HFD only (Fig. 1D_iii_). Although fat measures were impacted in female F_1_ and F_2_ HFD, they did not exhibit glucose impairment (Fig. 1D_iv-v_). Analysis of the area under the curve (AUC) for the GTT was consistent with the male and female glycemic responses shown in the glucose curves (Fig. S2B_i-v_). In line with the observed glycemic responses, the insulin tolerance test and the corresponding AUC demonstrated that male F_0_ WT HFD and TG HFD were insulin insensitive (Fig. 1E_i_ and S2C_i_). Analysis of the AUC indicated that F_1_ WT HFD and F_1_ TG HFD were insulin insensitive (Fig. S2C_ii_). Like the glucose tolerance test, there were more pronounced impairments revealed by the ITT for the F_1_ TG HFD in comparison to the F_1_ WT HFD and only the F_2_ TG HFD showed impaired insulin sensitivity (Fig. 1E_iii_ and Fig. S2C_ii-iii_). Like the GTT, there was no indication of insulin impairment in female HFD F_1_ nor F_2_ (Fig. 1E_iv-v_ and Fig. S2C_iv-v_).

To summarize, the effects of paternal high-fat diet on glucose homeostasis were sex-specific; male descendants had impaired glucose homeostasis, whereas females did not. Taken together, the assessments of weight and metabolic testing indicate that the TG descendants had enhanced responses to paternal obesity in comparison to WT descendants.

### 3.2 Paternal obesity was associated with altered liver gene expression in the F_0_-F_1_ with unique genes being differentially expressed in KDM1A descendants (F_1_-F_2_)

Obesity contributes to pathophysiological changes in gene expression in the liver [72]. To determine whether the altered metabolic status of HFD sires and their descendants (F_1_-F_2_) was associated with differential gene expression in the liver, we performed RNA-sequencing on the left lateral lobe (*lobus hepatis sinister lateralis*) of adult mice (F_0_-F_2_). Sequencing quality was high with RNA profiles having a Pearson correlation coefficient > 0.8 (Fig. S3A). Interestingly, principal component analysis of sequencing data revealed distinct hepatic transcriptomic profiles between males and females that was independent of experimental group and genotype (Fig. S3C). We compared hepatic transcriptome profiles by diet, sex, genotype and generation using a gene-level analysis at single-transcript resolution [55]. As expected, obesity was associated with differential liver gene expression. Liver from obese F_0_ WT males showed differential expression of 2,136 genes in comparison to non-obese F_0_ WT males (Fig. 2A, Lancaster p<0.05). Similarly, when comparing obese F_0_ TG to non-obese F_0_ TG, 1,476 genes were differentially expressed (Fig. 2B, Lancaster p<0.05). Of these differentially expressed genes (DEGs), 448 were commonly altered by obesity in both the F_0_ WT and F_0_ TG (p<0.0001; Fig. 2_i_). To identify which genes were altered due to genotype, we compared WT obese to TG obese and identified 524 DEGs, suggesting that obesity had a unique effect in TG mice due to an interaction between diet and genotype (Fig. 2C, Lancaster p<0.05).

**Figure 2:**
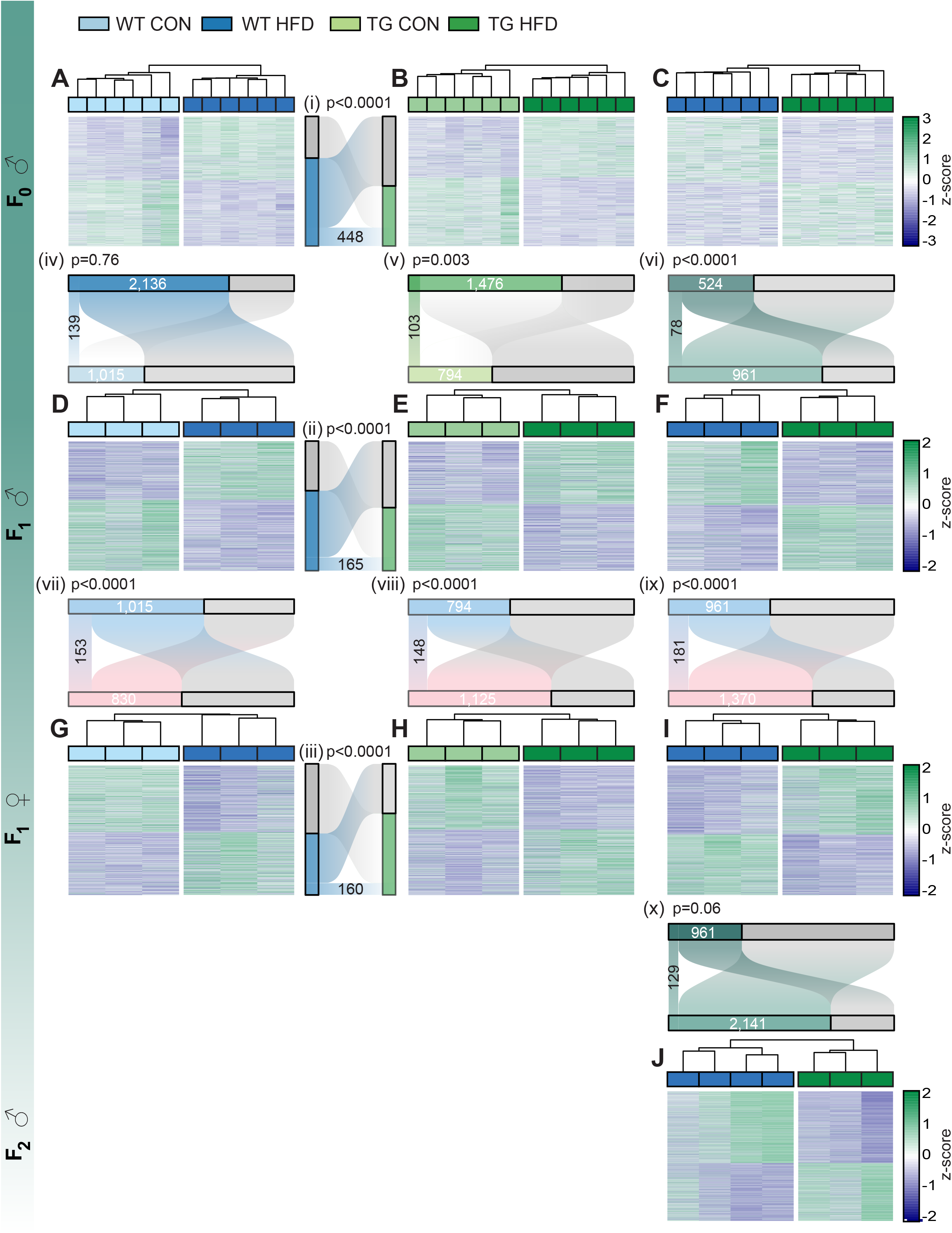
Paternal obesity is associated with altered gene expression in the livers of the F_0_-F_2_. A-J) Heatmaps of normalized expression values scaled by row (z-score) for transcripts that code for differentially expressed hepatic genes (Lancaster p-value<0.05) for each comparison assessed across sex and generation. Individual transcripts (rows) are ordered by k-means clustering and samples (columns) are arranged by hierarchical clustering, using complete-linkage clustering based on Euclidean distance. F_0_ WT CON vs WT HFD males (A), F_0_ TG CON vs TG HFD males (B), F_0_ WT HFD vs TG HFD males (C), F_1_ WT CON vs WT HFD males (D), F_1_ TG CON vs TG HFD males (E), F_1_ WT HFD vs TG HFD males (F), F_1_ WT CON vs WT HFD females (G), F_1_ TG CON vs TG HFD females (H), F_1_ WT HFD vs TG HFD females (I), and F_2_ WT HFD vs TG HFD males (J). i-x) Alluvial plots depicting frequency distributions of significant (colored boxes) and non-significant (grey boxes) genes for each comparison and their overlap across genotype (i-iii), across F_0_ and F_1_ males (iv-vi), across F_1_ males and females (vii-ix) and across F_1_ and F_2_ males (x). Significance of overlap between differentially expressed genes lists was calculated by Fisher’s exact test. P-values are included for each comparison above the respective alluvial plot.

To determine if the effects of paternal obesity on liver function were intergenerational, we compared the liver transcriptome of male and female F_1_. In comparison to F_1_ WT CON and TG CON males, livers of F_1_ WT HFD and TG HFD, showed differential expression of 1,015 and 794 genes (Fig. 2D and Fig. 2E, respectively, Lancaster p<0.05). A total of 165 DEGs overlapped between F_1_ WT and TG (p<0.0001; Fig. 2_ii_). Of the DEGs between the WT CON and HFD in the F_1_, 139 were the same deregulated genes as identified in the F_0_ WT CON vs HFD males (p=0.76; Fig 2_iv_). Similarly, there were 103 shared transcripts identified as differentially expressed between the F_1_ TG CON vs HFD, that were also altered in the F_0_ TG CON vs HFD (p=0.003; Fig 2_v_). This suggests that a common set of genes maintain dysfunction as a consequence of direct exposures to obesity and these changes are maintained in the F_1_ despite being fed a regular diet. When comparing genes altered by genotype in the F_1_ (WT HFD vs TG HFD), 961 were significantly altered (Fig 2F, Lancaster p<0.05), with 78 overlapping DEGs between the F_0_ and the F_1_ (p<0.0001; Fig 2_vi_). Demonstrating intergenerational (F_0_-F_1_) inheritance of metabolic dysfunction at the level of the liver, the metabolic regulators *Btg1* [73], *Cd300lg* [74], *FoxP4* [75], and *E4f1* [76], were differentially expressed in the livers of the obese F_0_ and their WT descendants. The overlap in deregulated genes between the F_0_ and F_1_ indicates that the metabolic phenotypes generated by the paternal HFD persist intergenerationally despite the F_1_ being fed a regular chow diet.

The last comparisons in liver transcriptomes were between the F_1_ male and female. Despite the female F_1_ having no metabolic phenotype detected by our measures, there was significantly altered gene expression in the livers of F_1_ female offspring of WT HFD vs WT CON sires (830; Fig 2G, Lancaster p<0.05). Of these, 153 were in common with the F_1_ male WT HFD sired offspring (p<0.0001; Fig 2_vii_). Likewise, the F_1_ female sired by TG HFD had 1,125 DEGs in comparison to females sired by TG CON (Fig. 2H, Lancaster p<0.05) with 148 in common with F_1_ male TG HFD sired offspring (p<0.0001; Fig. 2_viii_). Of these altered transcripts, 160 were in common between F_1_ female descendants of WT HFD and TG HFD (p<0.0001; Fig. 2_iii_). Like the F_1_ male TG HFD offspring, there were unique transcripts altered in F_1_ female TG HFD offspring (1,370; Fig. 2I, Lancaster p<0.05), with 181 differentially expressed in both F_1_ males and females (p<0.0001; Fig. 2_ix_). These may reflect genes impacted by genotype regardless of sex. An interesting finding from the F_2_ phenotyping was those transgenerational metabolic effects of the HFD were only detected in the male descendants of TG. Therefore, we only profiled F_2_ male livers by RNA-seq. This analysis revealed differential expression of 2,141 genes between the F_2_ WT HFD and TG HFD (Fig. 2J, Lancaster p<0.05) with 129 overlapping with the F_1_ WT HFD vs TG HFD males (p=0.06; Fig 2_x_). We identified 12 genes that showed transgenerational deregulated expression across the F_0_-F_2_, (WT HFD vs TG HFD), including *Eno3* which has been implicated in glycogen storage [77, 78], *Med23* which regulates insulin responsiveness [79], and *Prmt1* an epigenetic regulator implicated in liver glucose metabolism [80–82]. The number of differentially expressed genes increased every generation in comparisons between the WT HFD and the TG HFD (F_0_=524, F_1_=961, F_2_ = 2,141). This sustained deregulated gene expression in the livers of TG HFD F_2_, matches the enhanced metabolic phenotypes observed in only F_2_ TG HFD males, but not in the F_1_ WT HFD.

### 3.3 Paternal diet-induced obesity disrupts gene expression in functional processes that differ between genotypes, sexes and generations

To gain insight into the physiological implications of obesity-induced altered hepatic transcriptomes, we used a gene ontology (GO) approach combined with functional similarity clustering to compare processes in the liver impacted by diet across genotype and sex, and those impacted by genotype across generation (Fig. 3A-C, Supplemental files 1-3 and Table S5-7) [61]. Interactive heatmaps that facilitate in-depth probing of the gene frequency and the -log_10_ p-value of enriched GO terms within each cluster are found in Supplemental files 1-3. The non-interactive heatmaps are shown in Fig. 3. Overall, there were similar processes altered by obesity in F_0_ WT and TG livers, including lipid, amino acid, and small molecule metabolism (Fig. 3A, Supplemental file 1 and Table S5; clusters 1-5), homeostasis and environmental responses (clusters 8-10), and cellular differentiation and signalling (clusters 11-13). However, the gene frequency (# of genes annotated to that process) within processes differed by genotype.

**Figure 3:**
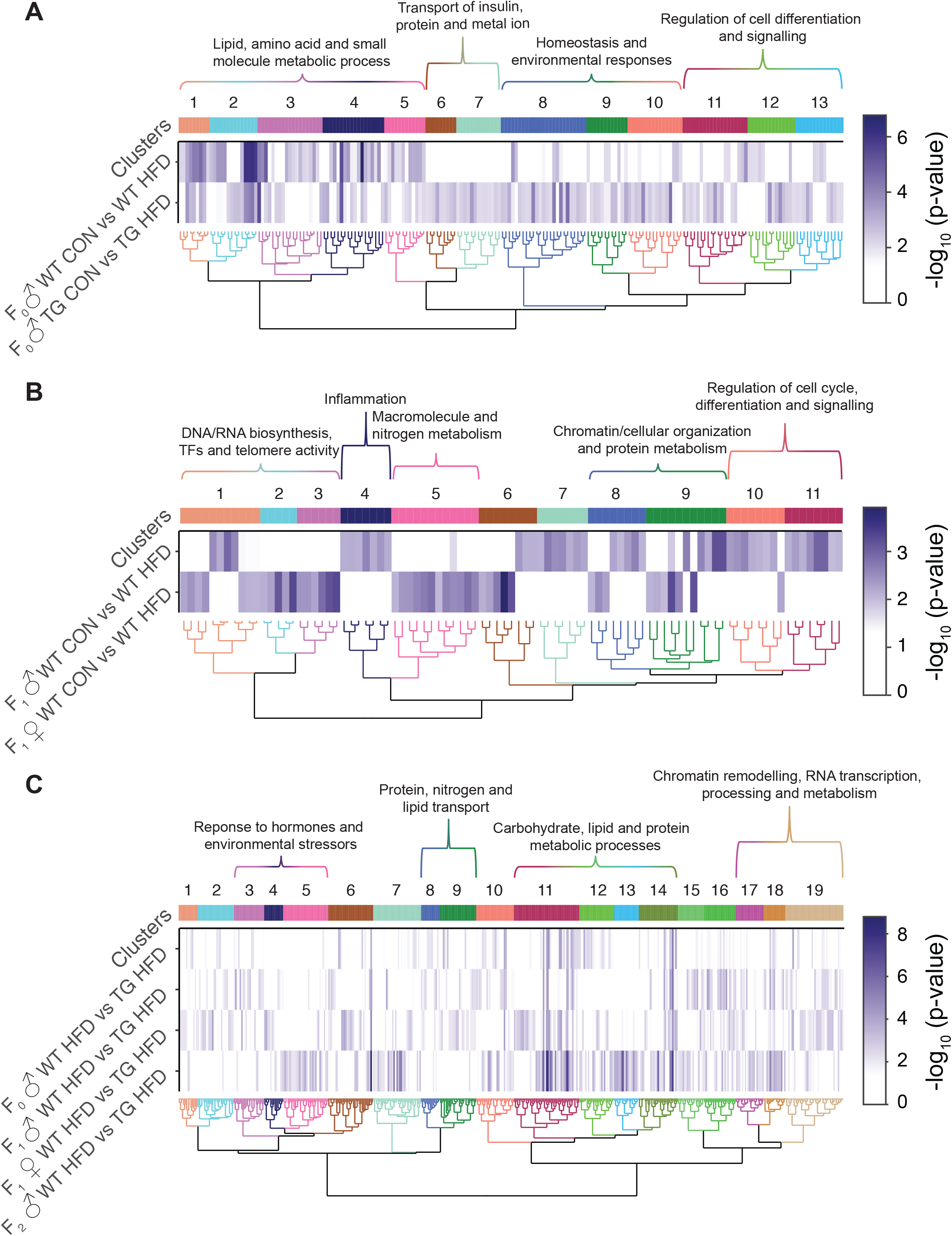
Obesity-induced hepatic transcriptome disturbances show functional similarities across genotype, sex and generation. A-C) Heatmaps of significant gene ontology (GO) terms clustered by functional similarity, comparing enriched biological functions for each comparison of interest across genotype (A), sex (B) and generation (C). Columns represent enriched GO terms which are ordered by hierarchical clustering based on Wang’s semantic similarity distance and *ward.D2* aggregation criterion. Each row represents a comparison of interest for which enriched GO terms were annotated based on the list of significant genes. The color gradient depicts the GO term enrichment significance (-log_10_ p-value). Interactive versions of these figures can be found in Supplemental files 1-3 and the complete lists of significantly enriched GO terms can be found in Tables S5-7.

When the altered functional pathways in F_1_ WT CON vs WT HFD were compared between males and females, there were clear impacts of paternal obesity on the liver biological pathways of offspring, and these differed by sex (Fig. 3B, Supplemental file 2 and Table S6). Reflecting sex differences, a greater number of GO terms related to inflammation (cluster 4), and cell cycle, differentiation and signalling regulation (clusters 10-11) were significantly enriched in males compared to females. Of note, genes involved in the regulation of proinflammatory cytokines were particularly enriched in males but not females (clusters 4). This concurs with the more severe phenotypes observed in the males. Conversely, genes involved in DNA/RNA biosynthesis, transcription factors and telomere activity (clusters 1-3), and macromolecule and nitrogen metabolism (cluster 5) were more enriched in females. Interestingly, pathways associated with chromatin and cellular organization and protein metabolism (clusters 8-9) were enriched by paternal obesity in both sexes.

Next, we compared the intergenerational and transgenerational effect of the interaction between the KDM1A transgene with obesity in terms of differences in process enrichment across generations when comparing F_0-2_ WT HFD with F_0-2_ TG HFD (Fig. 3C, Supplemental file 3 and Table S7). Reflecting the increasing generational changes in liver gene expression in the TG HFD descendants, there was an increase in the number of significantly enriched GO terms when comparing WT HFD vs TG HFD across generations (F_0_ male=79; F_1_ male=118; F_1_ female=159; F_2_ male=206; Supplemental file 4). This finding is concordant with the metabolic phenotypes detected in the TG HFD male F_2_ descendants, but not in the F_2_ male WT HFD (Fig. 1, Fig. S1, Fig. S2; Fig. 3C and Supplemental file 3). There was an enrichment in the differential expression of genes with functions related to inflammation and environmental response (clusters 3-5), metabolic processes (clusters 11-14), and chromatin remodelling and transcription (clusters 17-19). These enriched pathways in hepatic differentially expressed genes might reflect the interaction between obesity and the KDM1A transgene in the F_0_ sperm associated with the uniquely more severe and transgenerational phenotypes in TG HFD descendants (Fig. 3C).

### 3.4 Obesity in combination with germline expression of the KDM1A transgene increases differential enrichment of sperm H3K4me3 at genes involved in metabolism and development

We hypothesized that the sperm epigenome at the level of H3K4me3 would be altered by obesity and that this effect would be enhanced in KDM1A TG males with pre-existing alterations in sperm H3K4me3. To test these hypotheses, we performed native chromatin immunoprecipitation followed by deep sequencing (ChIP-seq) targeting histone H3K4me3, using sperm from individual WT or TG males fed either a CON or HFD (N=5 per experimental group, on average 33.3 million reads per sample with an alignment rate of 97%; Table S8). H3K4me3 localized to 30,745 genomic regions, with a Spearman correlation coefficient of 0.98 between samples (Fig. 4A and Fig. S4). Principal component analysis of H3K4me3 profiles revealed a clear separation of samples according to dietary treatment within genotype groups (Fig. 4B-C). WT samples separated along Principal Component 1 (PC1) with 37.41% of variance attributed to diet (Fig. 4B; PERMANOVA, permutation-based p=0.01). TG samples separated on PC1 with 32.68% of the variability, with diet as the second source of variance (PC2), at 25.56% (Fig. 4C; PERMANOVA, permutation-based p = 0.009).

**Figure 4:**
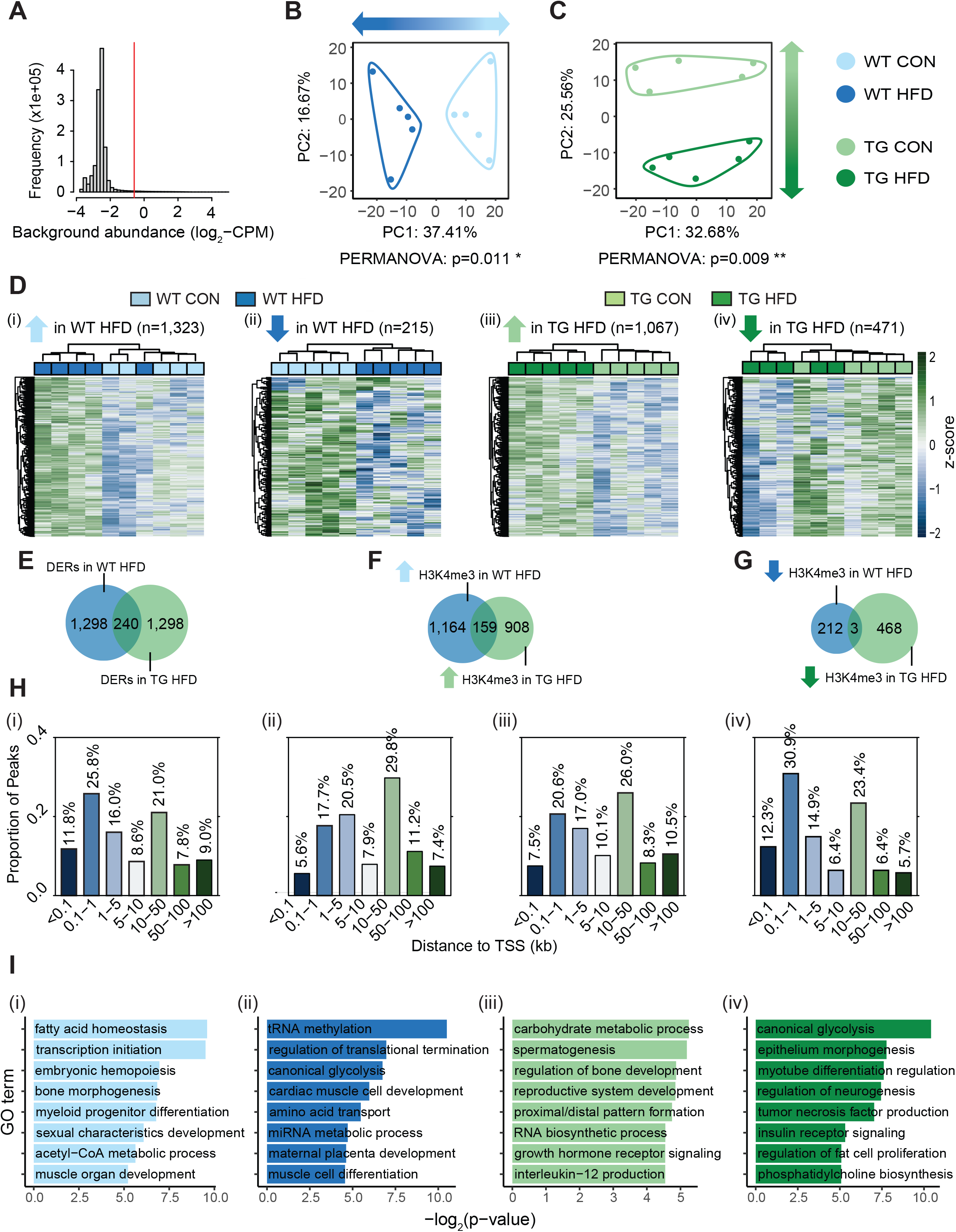
Genomic location, directionality change and functions of regions with altered H3K4me3 enrichment by obesity. A) Histogram showing frequency distributions of read abundances in 150 bp windows throughout the genome. Windows with an abundance below log_2_(4) fold over background bins of 2,000 bp were filtered out as indicated by the vertical red line. Enriched regions less than 100 bp apart were merged for a maximum width of 5,000 bp, conferring a total of 30,745 merged enriched regions. Reads were counted in merged enriched regions and normalized counts were used for downstream analyses. (see Material and Methods) B-C) Principal component analysis on normalized counts at merged enriched regions comparing WT CON vs WT HFD (B) and TG CON vs TG HFD (C). The top 5% regions contributing to separation of samples along Principal Component 1 (in B; PC1; x axis) or PC2 (in C; y axis) were selected. The PERMANOVA p-values indicating significance associated with dietary treatment are included under each PCA plot. D) Heatmaps of log2 normalized counts of deH3K4me3 regions in sperm with increased enrichment in WT HFD (i; n=1,323), decreased enrichment in WT HFD (ii; n=215), increased enrichment in TG HFD (iii; n=1,067) and decreased enrichment in TG HFD (iv; n=471) in each group. Samples (columns) and regions (rows) are arranged by hierarchical clustering using complete-linkage clustering based on Euclidean distance. Colored boxes indicate sample groups (light blue=WT CON, dark blue=WT HFD, light green=TG CON, dark green=TG HFD). E-G). Venn diagrams showing the overlap of deH3K4me3 in sperm of WT HFD (blue) and in TG HFD (green), for all detected regions (E), those gaining H3K4me3 (F) and those losing H3K4me3 (G). H) Barplots showing the distribution of altered regions based on the distance from the TSS of the nearest gene, for regions with increased enrichment in WT HFD (i; n=1,323), decreased enrichment in WT HFD (ii; n=215), increased enrichment in TG HFD (iii; n=1,067), and decreased enrichment in TG HFD (iv; n=471). The color gradient represents the distance of the regions to TSS in kilobase. I) Gene ontology analysis of diet-induced deH3K4me3 regions at promoters with increased enrichment in WT HFD (i; n=381), decreased enrichment in WT HFD (ii; n=34), increased enrichment in TG HFD (iii; n=230) and decreased enrichment in TG HFD (iv; n=150). Barplots show 8 selected significant GO terms with their respective -log_2_(p-value). Tables S9-12 include the complete lists of significantly enriched GO terms.

To focus our analysis on the regions most impacted by diet, we selected the top 5% differentially enriched H3K4me3 regions (deH3K4me3, n=1,538) in each genotype (PC1 in WT, PC2 in TG) (Fig. 4D_i-iv_). The genome distribution analysis for specific annotations showed that obesity-sensitive H3K4me3 regions were predominantly located in CpG islands, promoters, exons, and intergenic regions (Fig. S5). To a lesser extent, deH3K4me3 also occurred at transposable elements (LINE, SINE and LTR), where epigenetic de-repression is associated with the use of alternative promoters and long- and short-range enhancers that are implicated in embryo development and pluripotency [83] (Fig. S5). Representative genome browser tracks (Fig. S5) showing enrichment gains and losses for H3K4me3 at gene promoters are shown for *Pde1c* (phosphodiesterase 1C; affects the olfactory system), *Bcdin3d* (RNA methyltransferase; highly expressed in embryonic development), *Sh2d4a* (Sh2 domain containing protein 4A; expressed during development and associated with endocrine and liver function), and *Col15a1* (collagen Type XV alpha 1; involved in cell differentiation and various system development) [84].

Next, we compared the regions of H3K4me3 that were altered by obesity, their genomic location, directionality change and functions between diets and genotype (Fig. 4). As a response to obesity, H3K4me3 enrichment gains were more predominant than losses for both F_0_ WT HFD and TG HFD (Fig. 4D). In the WT HFD, 1,323 regions gained and 215 lost H3K4m3 in comparison to the WT CON (Fig. 4D_i-ii_). Similarly, in the F_0_ TG HFD sperm, 1,067 regions gained and 471 lost H3K4me3 in comparison to the F_0_ TG CON (Fig 4D_iii-iv_). Regions with deH3K4me3 in WT HFD had an 15.6% overlap (240/1,538 regions) with those of TG HFD (Fig. 4E). Of those common 240 regions, 162 had the same directionality change in both WT and TG HFD, with 159 regions with a gain and 3 regions with a reduction in H3K4me3 enrichment (Fig. 4F and Fig. 4G, respectively). The non-overlapping regions of deH3K4me3 in WT HFD and TG HFD sperm could be a consequence of genetic-epigenetic interactions where the TG mice respond uniquely to obesity as was observed in the phenotypic characterization. The proximity to the transcription start site (TSS) of the deH3K4me3 regions in sperm altered by obesity in the F_0_ WT HFD and TG HFD were similar (Fig. 4H).

Next, we performed a GO enrichment analysis on promoters to gain functional insight into the genes with obesity-responsive changes in sperm H3K4me3 enrichment and how they may relate to the developmental origin of offspring phenotypes. Notably, deH3K4me3 genes were identified in processes related to metabolism, inflammatory processes, and one-carbon cycle metabolism (Fig. 4I_i-iv_; Tables S7-10). Some of the significantly enriched pathways are concordant with disturbed metabolic phenotypes of the F_0_-F_2_ including for example, carbohydrate metabolic processes, glycolysis, growth hormone signaling and insulin signaling (Fig 4I, Tables S7-10).

The metabolic phenotypes of WT HFD and TG HFD descendants were similar, although the F_1-2_ TG HFD showed enhanced metabolic abnormalities. We hypothesized that these enhanced metabolic disturbances may relate to the greater degree of H3K4me3 alteration in F_0_ sperm, the directionality of the change (gain versus loss), and the functionality of genes bearing alterations. Together these factors could lead to increased disturbances of embryonic metabolic gene expression and more profound adult disease. Interestingly, when comparing WT CON with TG HFD sperm, samples separated along PC2, with 26.69% of variance associated with genotype and diet (Fig. 5A; PERMANOVA, permutation-based p=0.006). Of the top 5% impacted regions selected (n=1,538), a greater proportion showed a gain of enrichment for H3K4me3 in TG HFD sperm in comparison to WT CON (Fig. 5B, n=1,071 regions with gains; Fig. 5C. n=467 regions with losses). We analyzed the detected regions impacted by genotype and diet (n=1,538) for differential enrichment to determine whether obesity in combination with KDM1A overexpression led to greater changes in H3K4me3 enrichment. This analysis identified 264 regions with a significant linear trend, where TG HFD sperm showed a greater degree of change in enrichment, and TG CON and WT HFD showed intermediate changes in comparison to WT CON (Fig. 5D-E, adjusted p<0.2). There were only 9 significant regions with further increase in H3K4me3 in the TG HFD (Fig. 5D), whereas 255 regions showed a greater loss of H3K4me3 enrichment in the TG HFD (Fig. 5E). Consistent with the stronger metabolic phenotypes observed in the TG HFD F_1-2_, the functional analysis of the promoters showing significant linear trends (n=104) for H3K4me3 across experimental groups occurred at genes implicated in metabolic and cardiovascular disease progression (Fig. 5F, Table S13).

**Figure 5:**
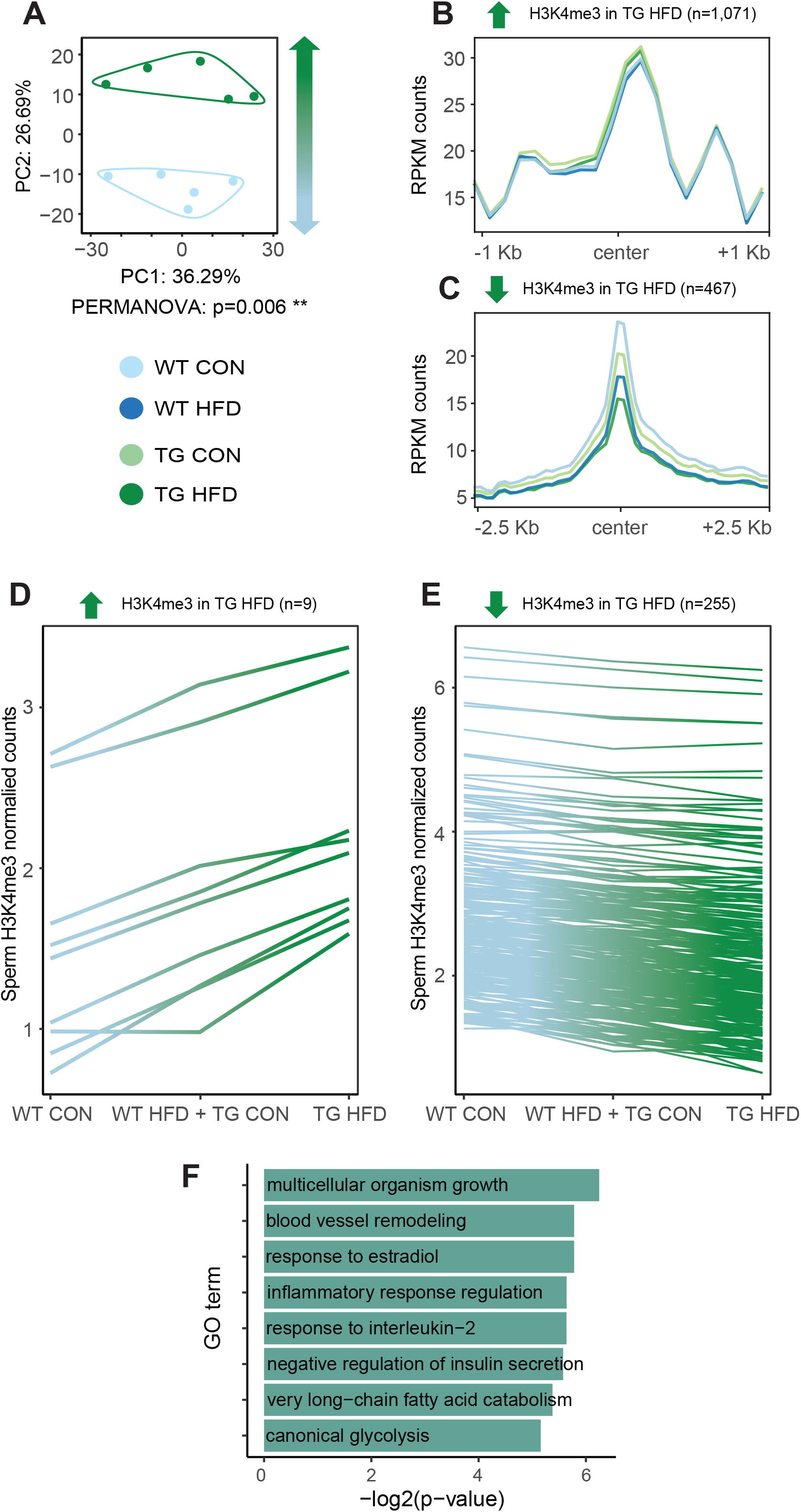
Additive effects of KDM1A overexpression and diet-induced obesity in the sperm epigenome at the level of H3K4me3. A) Principal component analysis on normalized counts at merged enriched regions comparing WT CON vs TG HFD. The top 5% regions contributing to separation of samples along Principal Component 2 (PC2; y axis) were selected. The PERMANOVA p-value under the plot indicates significance. B-C) Profile plots of RPKM H3K4me3 counts +/- 1 kilobase around the center of regions with increased H3K4me3 (B) and +/- 2.5 kilobase around the center of regions with decreased H3K4me3 enrichment in TG HFD (C). D-E) Line plots showing the median of normalized sperm H3K4me3 counts for each experimental group at regions showing a significant trend (n=264, adjusted p-value<0.2) with a linear increase in H3K4me3 enrichment (D; n=9) or a linear decrease in H3K4me3 enrichment (E; n=255) from WT CON, WT HFD, TG CON to TG HFD groups. F) Gene ontology analysis on the regions associated with a significant linear trend at promoters (n=104). Barplots show 8 selected significant GO terms with their respective -log_2_(p-value). Table S13 includes the complete list of significantly enriched GO terms.

### 3.5 Paternal obesity impacts sperm H3K4me3 at regions that coincide with open chromatin and gene expression in pre-implantation embryos

We recently demonstrated that sperm H3K4me3 is transmitted to the embryo and associated with gene expression [9]. We hypothesized that obesity-altered sperm H3K4me3 is transmitted and associated with chromatin accessibility in the early embryo, which in turn could influence gene expression and offspring phenotypes. To assess this possibility, we investigated the relationship between deH3K4me3 in sperm in relation to H3K4me3 in the embryo, the oocyte and open chromatin, and embryonic gene expression [40–43]. In line with a preferential paternal contribution of H3K4me3 to the 2-cell embryo, regions enriched for H3K4me3 in sperm, including those altered by obesity are not enriched in the oocyte (Fig 6A). There was a strong association between sperm H3K4me3, chromatin accessibility and embryonic gene expression at the 4-cell and morula stages (Fig. 6A-B and Fig. S6A_i_). Strikingly, sperm H3K4me3 including obesity-sensitive regions are associated with open chromatin in pre-implantation embryos (Fig. 6A-B).

**Figure 6:**
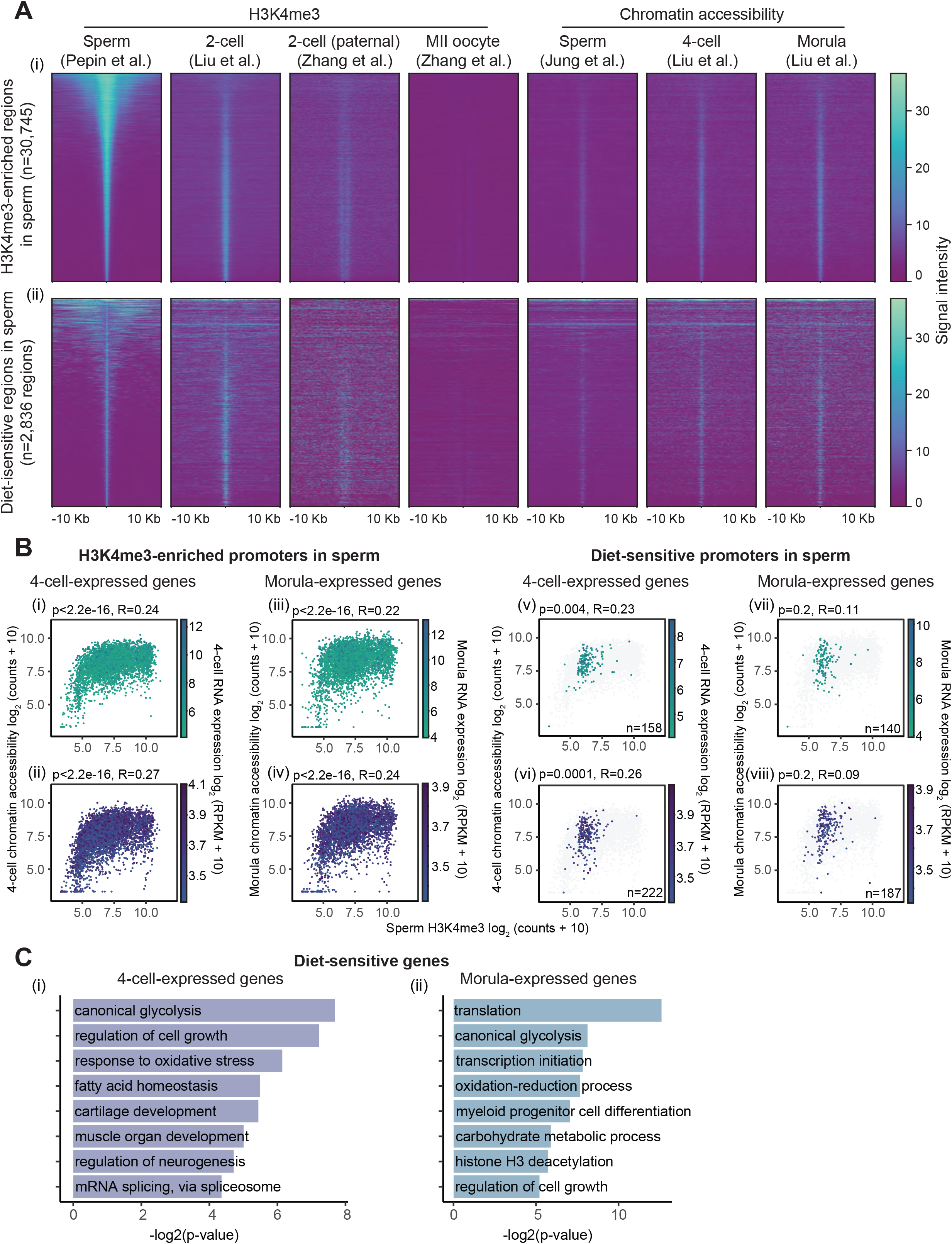
Sperm H3K4me3 regions sensitive to obesity occur at genes with an open chromatin state and expressed in the pre-implantation embryo. A) Heatmaps of RPKM counts signal +/- 10 kilobase around the center of regions enriched with H3K4me3 in sperm (i; n=30,745) and regions with obesity-induced deH3K4me3 in sperm (ii; n=2,836) for H3K4me3 enrichment levels in sperm (this study), 2-cell embryo (Liu *et al.*, 2016), 2-cell embryo on the paternal allele and MII oocyte (Zhang *et al.*, 2016), and for chromatin accessibility signal in sperm (Jung *et al.*, 2017), 4-cell embryo and morula embryo (Liu *et al.*, 2019). B) Scatterplots showing H3K4me3 enrichment in sperm (x axis; log2 counts + 10), chromatin accessibility signal (y axis; log2 counts + 10; (Jung *et al.*, 2017)) and gene expression levels (color gradient; log2 FPKM + 10; (Liu *et al.*, 2019)) in 4-cell (i,ii,v,vi) or in morula (iii,iv,vii,viii) embryos, at either all genes with promoters enriched with H3K4me3 in sperm (i-iv) or at diet-sensitive genes (v-viii). The top row of scatterplots includes lowly-expressed genes (bottom 50%) in 4-cell (i and v) or morula (iii or vii) embryos. The bottom row of scatterplots includes highly-expressed genes (top 50%) in 4-cell (ii and iv) or morula (vi and viii) embryos. Pearson’s correlation coefficients and their associated p-values are indicated above each scatterplot, comparing H3K4me3 enrichment in sperm versus H3K4me3 enrichment in 4-cell or morula embryos. C) Gene ontology analysis of genes expressed in the 4-cell (i) or the morula (ii) embryos, overlapping with diet-sensitive promoters in sperm. Barplots show 8 selected significant GO terms with their respective -log_2_(p-value). Tables S14-15 include the complete lists of significantly enriched GO terms.

To determine the functional relationship between the H3K4me3 obesity-altered regions and embryonic gene expression, we compared these with 4-cell and morula expressed genes and performed a gene ontology analysis. Of the sperm deH3K4me3 regions overlapping promoters (n=738), 51.8% (n=382) are expressed in the 4-cell embryos, 44.3% (n=327) are expressed in the morula embryos, and 39.7% (292) overlap in both (Fig. S6A_ii_). To gain insight into what obesity-altered H3K4me3 associated genes in sperm relate to embryonic gene expression, we performed a GO analysis on the promoters that are deH3K4me3 in sperm and the corresponding genes expressed in 4-cell and morula embryos (Fig. 6C_i-ii_). Again, supporting a role for sperm H3K4me3 in paternal transmission of metabolic disease, with both the 4-cell and the morula gene processes significantly enriched specific to metabolism (Fig. 6C_i-ii_ and Tables S12-13). Taken together these findings suggest a preferential contribution of H3K4me3 on the paternal chromatin in the early embryo that includes obesity-sensitive regions that may be instructive of metabolic-associated gene expression and a direct route for epigenetic inheritance.

### 3.6 HFD alters the sperm epigenome at regions instructive for placenta development

The placenta is a key extra-embryonic organ that represents the uterine-fetal interface and plays a central role in energy allocation, nutrient exchange, and developmental progression. Placental abnormalities have been linked to late onset cardiometabolic diseases, highlighting the importance of the *in utero* environment for metabolic health in adulthood [85]. Our gene ontology analysis on diet-induced deH3K4me3 regions in sperm revealed significant enrichment of genes involved in placenta development (Fig. 4I and Tables S7-10). Given the sperm epigenome influences placental gene expression [86], we were interested in the prospect that diet-induced epimutations in sperm affect placenta gene expression that could influence metabolic phenotypes across generations. To investigate this possibility, we compared the enrichment profiles of H3K4me3 in sperm, with H3K4me3 signal and gene expression data from trophectoderm (TE, the embryonic precursor of placental lineage) [40], and placenta [44, 87]. Most regions enriched with H3K4me3 in sperm showed strong H3K4me3 signal in TE and placenta (Fig. 7A), with 65.9% (n=8,663) and 79.4% (n=10,434) of H3K4me3-enriched sperm promoters (n=13,142) expressed in these tissues, respectively (Fig. S6B_i_). Of the 738 deH3K4me3 regions localizing to promoters in sperm, 56.8% (n=418) were expressed in the trophectoderm, 76.8% (n=567) were expressed in the placenta, and 54.6% (n=403) were expressed in both (Fig. S6B_ii_). Notably, gene ontology analysis of the shared H3K4me3 in sperm with TE and placenta revealed that there was an association with placenta function including at deH3K4me3 regions (Fig. 7B_i,iii_, Tables S14 and S16). The GO analysis of the sperm H3K4me3 regions that were not common with TE and placenta were involved in spermatogenesis, fertilization and sperm function (Fig. 7B_ii and iv_, Tables S15 and S17).

**Figure 7:**
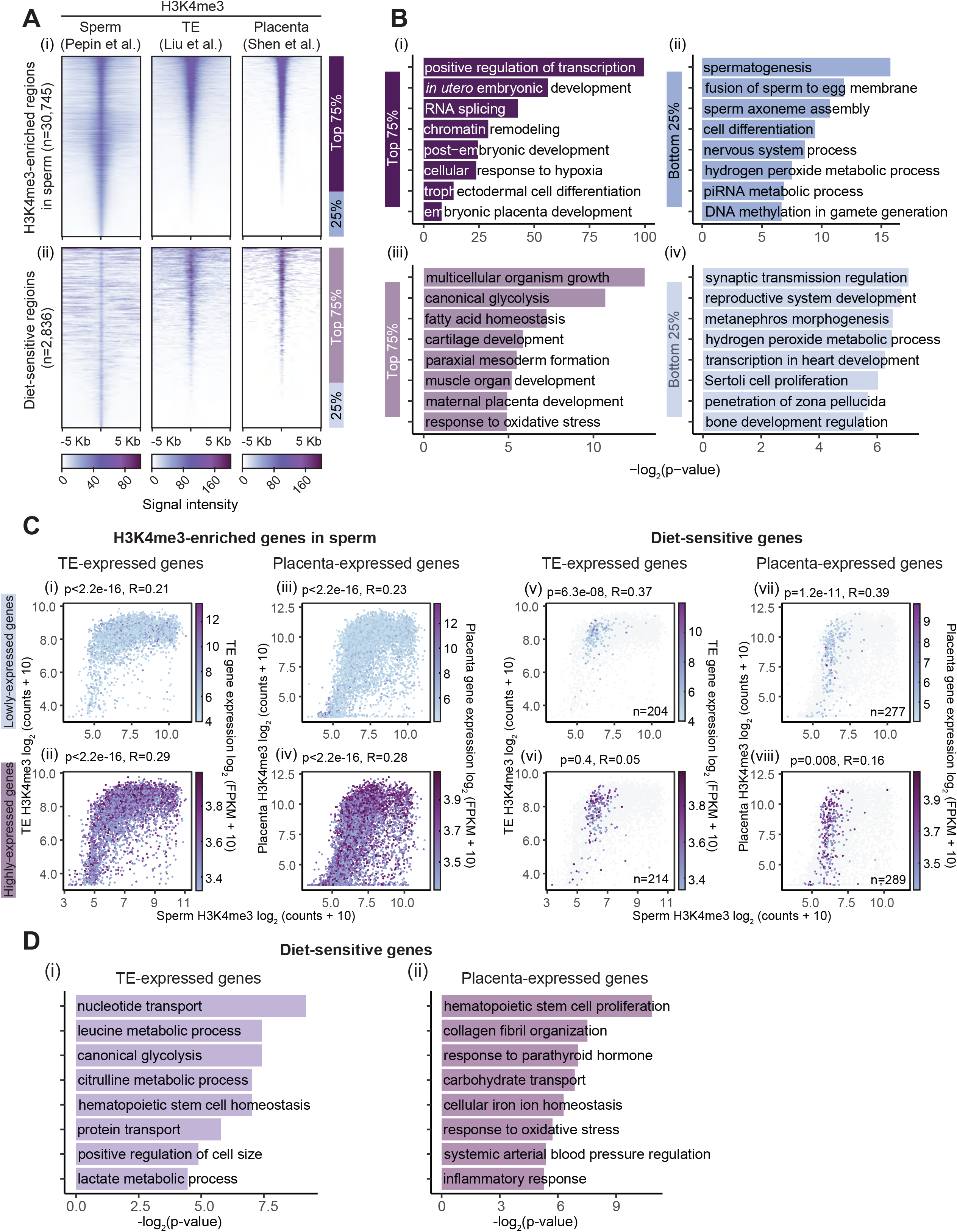
Obesity-induced deH3K4me3 regions overlap with genes marked by H3K4me3 and expressed in the trophectoderm and placenta. A) Heatmaps of RPKM counts signal +/- 5 kilobase around the center of regions enriched with H3K4me3 in sperm (i; n=30,745) and at regions with diet-induced deH3K4me3 in sperm (n=2,836) for H3K4me3 enrichment levels in sperm (this study), trophectoderm (TE) (Liu *et al.*, 2016) and placenta (Shen *et al.*, 2012). B) Gene ontology analysis of regions enriched with H3K4me3 in sperm, TE and placenta (top 75% from A i) (i), regions enriched with H3K4me3 in sperm only (bottom 25% from A i) (ii), diet-sensitive regions enriched with H3K4me3 in sperm, TE and placenta (top 75% from A ii) (iii), and diet-sensitive regions enriched with H3K4me3 in sperm only (bottom 25% from A ii) (iv). Barplots show 8 selected significant GO terms with their respective -log_2_(p-value). Tables S16-19 include the complete lists of significantly enriched GO terms. C) Scatterplots showing H3K4me3 enrichment at promoters in sperm (x axis; log2 counts + 10), H3K4me3 enrichment (y axis; log2 counts + 10) and gene expression levels (color gradient; log2 FPKM + 10) in the trophectoderm (i,ii,v,vi; (Liu *et al.*, 2016)) or in the placenta (iii,iv,vii,viii; (Shen *et al.*, 2012; Chu *et al.*, 2019)), at either all genes with promoters enriched with H3K4me3 in sperm (i-iv) or at diet-sensitive genes (v-viii). The top row of scatterplots includes lowly-expressed genes (bottom 50%) in trophectoderm (i and v) or placenta (iii or vii). The bottom row includes highly-expressed genes (top 50%) in trophectoderm (ii and iv) or placenta (vi and viii). Pearson’s correlation coefficients and associated p-values are indicated above each scatterplot, comparing H3K4me3 enrichment in sperm versus H3K4me3 enrichment in the trophectoderm or placenta. D) Gene ontology analysis of genes expressed in the trophectoderm (i) or the placenta (ii), overlapping with diet-sensitive promoters in sperm. Barplots show 8 selected significant GO terms with their respective -log_2_(p-value). Tables S20-21 include the complete lists of significantly enriched GO terms.

Next, we compared gene enrichment of sperm H3K4me3 with low- and high-expressed genes in the TE and placenta. Suggesting an influential role of sperm H3K4me3, the highly expressed genes and to a lesser extent the lowly expressed genes in TE and placenta were positively correlated with sperm H3K4me3 (Fig. 7C_i-iv_). Notably when the same comparisons were made with the deH3K4me3 there was a significant relationship with both lowly- and highly-expressed placenta genes (p=1.2e-11 and p=0.008, respectively; Fig. 7C_v-viii_). These included genes implicated in altered placenta hormonal profiles and preeclampsia such as *Ldoc1*, *Dab2ip*, and *Rgs2* [88–91]. In addition, the GO analysis of TE- and placenta-expressed genes that overlap with deH3K4me3 promoters are in line with the metabolic phenotypes in offspring (Fig. 7D_i-ii_, Tables S18-19). Taken together this analysis raises the possibility that obesity-induced alterations in sperm may influence embryonic and placenta gene expression to alter metabolic function of offspring.

### 3.7 Obesity-induced sperm epigenomic and hepatic transcriptomic alterations are unrelated

In a recent study, paternal low-protein diet was associated with reduced H3K9me2 at genes in sperm and were suggested to modulate gene expression profiles in the liver [92]. We aimed to assess whether a similar association between obesity-induced deH3K4me3 in sperm would relate to differential expression in the livers of the next generation. We focused on the obesity-associated sperm deH3K4me3 at promoters in F_0_ sires and their relationship to differentially expressed genes in the liver (DEGs) of F_1_ males. This analysis revealed that genes with differential expression in livers (n=1,644) were by and large unrelated to genes bearing deH3K4me3 in sperm. Only 9.1% (n=67) of promoters with deH3K4me3 in sperm were differentially expressed in the liver of F_1_ males sired by HFD-fed sires (Fig. S7A-B). We then asked if deH3K4me3 promoters in sperm and liver DEGs had related biological functions. Strikingly, sperm- and liver-altered genes showed few functional similarities (Fig. S7C, Supplemental file 5 and Table S22). Functional pathways specifically enriched in deH3K4me3 promoters in sperm involved development and differentiation processes (clusters 12-15). As expected in a paternal obesity model, gene processes altered in offspring livers included: regulation of transcription and RNA splicing (clusters 1-3), protein and histone post-translational modifications (clusters 4-5), and metabolism of lipid, nitrogen and glucose (clusters 6-8). Pathways enriched in both the deH3K4me3 promoters in sperm and the DEGs in liver were involved in cell cycle, transport and signaling (clusters 16-19), and response to stress and inflammation (clusters 20-22). These commonly enriched pathways might reflect obesity-associated systemic inflammation which could affect multiple organs in a similar manner. These findings indicate that paternal obesity alters the sperm epigenome at distinct genes and functional pathways than those differentially expressed in offspring livers and fits with a developmental origin of adult metabolic dysfunction that could be related to alterations in gene expression in the embryo and placenta.

## 4 Discussion

In mammals, spermatogenesis is a highly complex cell differentiation process involving unique testis-specific gene expression programs that are accompanied by dynamic remodeling of the chromatin [93–96]. During this process, most histones are replaced by protamines to facilitate DNA compaction [93]. Interestingly, 1% of sperm histones are retained in mice and 15% in men [97, 98]. Retained histones are conserved across species from mice to men and are found at the gene regulatory regions implicated in spermatogenesis, sperm function, embryo development, metabolism and routine cellular processes [98–100]. We have shown that in human and mouse sperm histone H3 lysine 4 dimethylation (H3K4me2) and trimethylation (H3K4me3) localize to genes involved in metabolism and development [9,24,101]. Since this tantalizing discovery we and others have suggested that histones in sperm may directly influence embryonic gene expression and contribute to the developmental origin of adult disease. The findings of this study support histones serving in this mechanism of disease inheritance.

In spermatogenesis there are dynamic changes to the sperm epigenome including histone methylation which is susceptible to alterations induced by changes in methyl donor availability [5, 9]. Diets high in fat alter epigenetic programming, likely through the alteration of cellular metabolism, which influences the availability of methyl donors and/or the activation or inactivation of chromatin modifying enzymes. In overweight and obese individuals, homocysteine is consistently elevated, and associated with reduced B12 and folate [102, 103]. It follows that the obesity-induced alterations in H3K4me3 we report here could be a consequence of an altered methyl donor pool. Intriguingly, the effects of obesity on the paternal epigenome were linked with the metabolic dysfunction in the F_1_ and F_2_ descendants; deH3K4me3 occurred at the promoters of genes involved in fertility, metabolism, and placenta processes. Indicative of paternal transmission of sperm altered H3K4me3 as a mediator of metabolic dysfunction was the strong relationship between deH3K4me3, an open chromatin state and gene expression in embryos and placenta. However, a limitation of the study is that we did not examine H3K4me3 in sperm from the F_1_ and thus whether H3K4me3 abnormalities in sperm persist in the subsequent generation is unknown. In this model of diet-induced transgenerational inheritance and in others, offspring phenotypes are likely the consequence of a complex interplay between chromatin, DNA methylation and non-coding RNA in sperm and embryos. For example, a paternal low protein diet has been shown to alter testicular germ cell activating transcription factor 7 (ATF7) binding, and this was associated with differential sperm H3K9me2 and small RNA content in spermatocytes [92]. Elucidating whether there are common molecular pathways mediating inter- and trans-generational impacts of paternal diet remains to be determined.

The enhanced metabolic abnormalities observed in the descendants of obese F_0_ TG revealed an increased susceptibility to metabolic disease in the TG line. An explanation for this response is that the F_0_ TG were descendants from a lineage with pre-existing alterations in the sperm epigenome due to the genetic modification causing KDM1A overexpression. This genetic stress in combination with the environmental challenge of the HFD resulted in a more severely altered sperm epigenome in comparison to the WT, with consequent enhanced offspring phenotypes. Admittedly speculative, these findings suggest that the higher incidence of poor health in at-risk populations may be attributed to generational exposures to poor diet that leads to an accumulation of sperm epigenome errors that escape reprogramming.

Notably, paternal obesity-induced transgenerational metabolic disturbances in offspring were only observed in descendants of obese TG males. The phenomenon of transgenerational inheritance has been most documented in genetic mouse models of epigenetic inheritance and studied in relation to DNA methylation patterns. These include the Avy agouti model [21,104,105], the kinky tail model (Axin^Fu^ allele) [106], and in mice bearing a mutation in the *Mtrr* gene, a folate metabolism enzyme [107]. In the context of environmental challenges, paternal transgenerational inheritance has been associated with altered sperm DNA methylation when there has been gestational exposure to toxicants and undernutrition [108, 109], and in a non-genetic pharmacologically-induced prediabetes model begun at weaning [110]. Taken together, this growing body of evidence indicates that transgenerational inheritance occurs under genetic influence, or when exposures coincide with developmental programming. The male F_0_ mice in this study were exposed to the paternal HFD from weaning and not *in utero*, which may account for why transgenerational effects were not observed in WT HFD descendants. Another possibility is that transgenerational responses in the WT may have become detectable in older mice.

Our analysis indicates that the inherited metabolic disturbances observed in adult descendants originated early in development. In rodent models, paternal obesity and *in utero* undernutrition has been linked to altered gene expression in offspring livers and pancreatic islets with some minor links to concordant DNA methylation changes [109–111]. It has been suggested that diet-associated alterations in DNA methylation in sperm are retained through embryogenesis and maintained in adult tissues mediating paternally-induced phenotypes [109, 110]. Consistent with these studies, altered hepatic gene expression occurred in F_1-2_ offspring of obese sires. In contrast, we observed minimal overlap of genes and functional pathways between altered H3K4me3 enrichment in sperm, with those differentially expressed in F_1_ livers. Instead, we demonstrate a significant overlap of obese sperm H3K4me3 profiles with the expression of metabolic-related genes in the embryo and placenta. Based on these findings, we suggest that the metabolic phenotypes we observe originate in early embryogenesis and through changes in placental gene expression.

There is a bounty of research linking maternal obesity to adverse metabolic consequences for the offspring that coincide with altered placental gene expression and function [112, 113]. On the other hand, it is an emerging concept that the paternal environment including factors such as diet and age can influence placental development and function. It is known that paternally expressed genes contribute to placental growth, trophoblast invasion and insulin resistance and adiposity [86,114–119]. In humans, errors in epigenomic programming have been associated with gestational trophoblast disease and pre-eclampsia, but the role of the obese father in these conditions has been entirely unexplored [120, 121]. Previous studies support a connection between paternal diets, obesity, and placental dysfunction as a developmental route to metabolic disease in children. For example, we have shown that a folate deficient paternal diet and altered sperm DNA methylation coincided with deregulated placenta gene expression of *Cav1* and *Txndc16* [5]. Moreover, paternal obesity in mice has been attributed to defective placental development [115,122,123]. In women, altered DNA methylation in the regulation of some genes in preeclampsia has been established. However, many genes with deregulated expression were not associated with DNA methylation raising the possibility of altered chromatin signatures leading to abnormal gene expression in this placental disorder [124]. Indeed, upregulated expression of LncRNA by increased H3K4me3 has been observed in preeclampsia placentas [125], and the levels of H3K4me3 as detected by immunohistochemistry are decreased [126]. Until now the connection between sperm chromatin and placenta function has been unexplored. Our analyses revealed that most of the obesity-altered H3K4me3 at promoters occurred at loci involved in placental development and inflammatory processes, with 56.6% and 76.8% of deH3K4me3 occurring at promoters expressed in the trophectoderm and placenta, respectively. Remarkably, deregulated expression of genes implicated in inflammation has been implicated in hypertensive disorders in pregnancy including pre-eclampsia. Hypertensive disorders in pregnancy have been associated with increased risk for developing cardiovascular disease [117]. This raises the possibility that the paternal sperm epigenome may influence maternal health during pregnancy in addition to that of the developing fetus.

As in previous studies we found that paternal obesity resulted in sex-specific differences in metabolism and fat accruement with males being more impacted. The underlying mechanisms that lead to the greater susceptibility of males may be related to sexually dimorphic placental gene expression [127]. In support of this possibility, paternal environment (diet) influenced placental function in a sex-specific manner [122]. Alternatively, different metabolic responses in male and female offspring may be due to hormonal responses where estrogen has been shown to protect against altered glucose homeostasis [29, 128].

In summary, we provide evidence that paternal obesity is associated with H3K4me3 signatures in sperm which could contribute to the inheritance of metabolic disease. In addition, we identified links between sperm regions bearing obesity-altered H3K4me3, with placenta and embryonic H3K4me3, and the regulation of gene expression in these tissues. Important next steps to better understand disease inheritance related to paternal obesity, sperm chromatin and placental function will be to explore this possibility using embryonic and placenta tissue from pregnancies sired by obese males. The translational validation of these findings will be important in developing intervention strategies focused on paternal factors that could impact the health of future generations [129].

## Acknowledgements

We thank the team at the McGill University Small Animal Research Unit, Génome Québec Innovation Centre for performing the sequencing, and Dr. Deborah Sloboda (McMaster University) for advice on metabolic phenotyping methods. This research was funded by the Canadian Institute of Health Research (CIHR) grants to SK (358654 and 350129).

## CRediT author statement

**Anne-Sophie Pépin:** Methodology, Data curation, Software, Investigation, Formal analysis, Visualization, Writing – Original draft preparation

**Christine Lafleur**: Resources, Investigation

**Romain Lambrot:** Resources

**Vanessa Dumeaux**: Software, Resources, Writing – Review & Editing

**Sarah Kimmins**: Supervision, Conceptualization, Funding acquisition, Writing – Original draft preparation

## References

1. Gernand, A.D., Schulze, K.J., Stewart, C.P., West, K.P.J., Christian, P., 2016. Micronutrient deficiencies in pregnancy worldwide: health effects and prevention. Nature Reviews. Endocrinology 12(5): 274–89, Doi: 10.1038/nrendo.2016.37.

2. Braun, J.M., Messerlian, C., Hauser, R., 2017. Fathers Matter: Why It’s Time to Consider the Impact of Paternal Environmental Exposures on Children’s Health. Current Epidemiology Reports 4(1): 46–55, Doi: 10.1007/s40471-017-0098-8.

3. Donkin, I., Barrès, R., 2018. Sperm epigenetics and influence of environmental factors. Molecular Metabolism 14: 1–11, Doi: https://doi.org/10.1016/j.molmet.2018.02.006.

4. Eberle, C., Kirchner, M.F., Herden, R., Stichling, S., 2020. Paternal metabolic and cardiovascular programming of their offspring: A systematic scoping review. PloS One 15(12): e0244826, Doi: 10.1371/journal.pone.0244826.

5. Lambrot, R., Xu, C., Saint-Phar, S., Chountalos, G., Cohen, T., Paquet, M., et al., 2013. Low paternal dietary folate alters the mouse sperm epigenome and is associated with negative pregnancy outcomes. Nature Communications 4, Doi: 10.1038/ncomms3889.

6. Radford, E.J., Ito, M., Shi, H., Corish, J.A., Yamazawa, K., Isganaitis, E., et al., 2014. In utero undernourishment perturbs the adult sperm methylome and intergenerational metabolism. Science 345(6198): 1255903, Doi: 10.1126/science.1255903.

7. Donkin, I., Versteyhe, S., Ingerslev, L.R., Qian, K., Mechta, M., Nordkap, L., et al., 2016. Obesity and bariatric surgery drive epigenetic variation of spermatozoa in humans. Cell Metabolism 23(2): 369–78, Doi: 10.1016/j.cmet.2015.11.004.

8. Wu, H., Estill, M.S., Shershebnev, A., Suvorov, A., Krawetz, S.A., Whitcomb, B.W., et al., 2017. Preconception urinary phthalate concentrations and sperm DNA methylation profiles among men undergoing IVF treatment: a cross-sectional study. Human Reproduction (Oxford, England) 32(11): 2159–69, Doi: 10.1093/humrep/dex283.

9. Lismer, A., Dumeaux, V., Lafleur, C., Lambrot, R., Amour, J.B., Lorincz, M.C., et al., 2021. Histone H3 lysine 4 trimethylation in sperm is transmitted to the embryo and associated with diet-induced phenotypes in the offspring Article Histone H3 lysine 4 trimethylation in sperm is transmitted to the embryo and associated with diet-induce. Developmental Cell: 1–16, Doi: 10.1016/j.devcel.2021.01.014.

10. Pilsner, J.R., Shershebnev, A., Wu, H., Marcho, C., Dribnokhodova, O., Shtratnikova, V., et al., 2021. Aging-induced changes in sperm DNA methylation are modified by low dose of perinatal flame retardants. Epigenomics 13(4): 285–97, Doi: 10.2217/epi-2020-0404.

11. Voight, B.F., Scott, L.J., Steinthorsdottir, V., Morris, A.P., Dina, C., Welch, R.P., et al., 2010. Twelve type 2 diabetes susceptibility loci identified through large-scale association analysis. Nature Genetics 42(7): 579–89, Doi: 10.1038/ng.609.

12. Kaati, G., Bygren, L.O., Edvinsson, S., 2002. Cardiovascular and diabetes mortality determined by nutrition during parents’ and grandparents’ slow growth period. European Journal of Human Genetics 10(11): 682–8, Doi: 10.1038/sj.ejhg.5200859.

13. Pembrey, M.E., Bygren, L.O., Kaati, G., Edvinsson, S., Northstone, K., Sjöström, M., et al., 2006. Sex-specific, male-line transgenerational responses in humans. European Journal of Human Genetics 14(2): 159–66, Doi: 10.1038/sj.ejhg.5201538.

14. Lumey, L.H., Poppel, F.W.A. Van., 2013. The Dutch Famine of 1944-45 as a Human Laboratory: Changes in the Early Life Environment and Adult Health. In: Lumey, L. H. and Vaiserman, A., editor. Early Life Nutrition and Adult Health and Development, Nova Science Publishers, Inc. p. 59–76.

15. Siklenka, K., Erkek, S., Godmann, M., Lambrot, R., McGraw, S., Lafleur, C., et al., 2015. Disruption of histone methylation in developing sperm impairs offspring health transgenerationally. Science 350(6261), Doi: 10.1126/science.aab2006.

16. Dalgaard, K., Landgraf, K., Heyne, S., Lempradl, A., Longinotto, J., Gossens, K., et al., 2016. Trim28 Haploinsufficiency Triggers Bi-stable Epigenetic Obesity. Cell 164(3): 353–64, Doi: 10.1016/j.cell.2015.12.025.

17. Miska, E.A., Ferguson-smith, A.C., 2016. Transgenerational inheritance: Models and mechanisms of non – DNA sequence – based inheritance 354(6308): 778–82.

18. Carone, B.R., Fauquier, L., Habib, N., Shea, J.M., Hart, C.E., Li, R., et al., 2010. Paternally induced transgenerational environmental reprogramming of metabolic gene expression in mammals. Cell 143(7): 1084–96, Doi: 10.1016/j.cell.2010.12.008.

19. Grandjean, V., Fourré, S., De Abreu, D.A.F., Derieppe, M.A., Remy, J.J., Rassoulzadegan, M., 2015. RNA-mediated paternal heredity of diet-induced obesity and metabolic disorders. Scientific Reports 5(June): 1–9, Doi: 10.1038/srep18193.

20. Chen, Q., Yan, M., Cao, Z., Li, X., Zhang, Y., Shi, J., et al., 2016. Sperm tsRNAs contribute to intergenerational inheritance of an acquired metabolic disorder. Science 351(6271): 397–400, Doi: 10.1126/science.aad7977.

21. Cropley, J.E., Eaton, S.A., Aiken, A., Young, P.E., Giannoulatou, E., Ho, J.W.K., et al., 2016. Male-lineage transmission of an acquired metabolic phenotype induced by grand-paternal obesity. Molecular Metabolism 5(8): 699–708, Doi: 10.1016/j.molmet.2016.06.008.

22. de Castro Barbosa, T., Ingerslev, L.R., Alm, P.S., Versteyhe, S., Massart, J., Rasmussen, M., et al., 2016. High-fat diet reprograms the epigenome of rat spermatozoa and transgenerationally affects metabolism of the offspring. Molecular Metabolism 5(3): 184–97, Doi: 10.1016/j.molmet.2015.12.002.

23. Sharma, U., Conine, C.C., Shea, J.M., Boskovic, A., Derr, A.G., Bing, Y., et al., 2016. Biogenesis and function of tRNA fragments during sperm maturation and fertilization in mammals 351(6271): 391–6, Doi: 10.1126/science.aad6780.Biogenesis.

24. Lambrot, R., Siklenka, K., Lafleur, C., Kimmins, S., 2019. The genomic distribution of histone H3K4me2 in spermatogonia is highly conserved in sperm. Biology of Reproduction 100(6): 1661–72, Doi: 10.1093/biolre/ioz055.

25. Lambrot, R., 2021. Whole genome sequencing of H3 lysine 4 tri-methylation and DNA methylation in human sperm reveals regions of overlap and exclusion linked to fertility, development and epigenetic inheritance. Cell Reports CELL-REPOR.

26. Lismer, A., Siklenka, K., Lafleur, C., Dumeaux, V., Kimmins, S., 2020. Sperm histone H3 lysine 4 trimethylation is altered in a genetic mouse model of transgenerational epigenetic inheritance. Nucleic Acids Research, Doi: 10.1093/nar/gkaa712.

27. Stringer, J.M., Forster, S.C., Qu, Z., Prokopuk, L., O’Bryan, M.K., Gardner, D.K., et al., 2018. Reduced PRC2 function alters male germline epigenetic programming and paternal inheritance. BMC Biology 16(1): 1–20, Doi: 10.1186/s12915-018-0569-5.

28. Lesch, B.J., Tothova, Z., Morgan, E.A., Liao, Z., Bronson, R.T., Ebert, B.L., et al., 2019. Intergenerational epigenetic inheritance of cancer susceptibility in mammals. ELife 8: 1–29, Doi: 10.7554/elife.39380.

29. Lainez, N.M., Jonak, C.R., Nair, M.G., Ethell, I.M., Wilson, E.H., Carson, M.J., et al., 2018. Diet-induced obesity elicits macrophage infiltration and reduction in spine density in the hypothalami of male but not female mice. Frontiers in Immunology 9(SEP): 1–16, Doi: 10.3389/fimmu.2018.01992.

30. Ayala, J.E., Samuel, V.T., Morton, G.J., Obici, S., Croniger, C.M., Shulman, G.I., et al., 2010. Standard operating procedures for describing and performing metabolic tests of glucose homeostasis in mice. DMM Disease Models and Mechanisms 3(9–10): 525–34, Doi: 10.1242/dmm.006239.

31. Hisano, M., Erkek, S., Dessus-Babus, S., Ramos, L., Stadler, M.B., Peters, A.H.F.M., 2013. Genome-wide chromatin analysis in mature mouse and human spermatozoa. Nature Protocols 8(12): 2449–70, Doi: 10.1038/nprot.2013.145.

32. Lismer, A., Lambrot, R., Lafleur, C., Dumeaux, V., Kimmins, S., 2021. ChIP-seq protocol for sperm cells and embryos to assess environmental impacts and epigenetic inheritance. STAR Protocols 2(2): 100602, Doi: https://doi.org/10.1016/j.xpro.2021.100602.

33. Krueger, F., 2015. Trim galore. A Wrapper Tool around Cutadapt and FastQC to Consistently Apply Quality and Adapter Trimming to FastQ Files 516: 517.

34. Kim, D., Langmead, B., Salzberg, S.L., 2015. HISAT: a fast spliced aligner with low memory requirements. Nature Methods 12(4): 357–60, Doi: 10.1038/nmeth.3317.

35. Li, H., Handsaker, B., Wysoker, A., Fennell, T., Ruan, J., Homer, N., et al., 2009. The Sequence Alignment/Map format and SAMtools. Bioinformatics (Oxford, England) 25(16): 2078–9, Doi: 10.1093/bioinformatics/btp352.

36. Pertea, M., Pertea, G.M., Antonescu, C.M., Chang, T.-C., Mendell, J.T., Salzberg, S.L., 2015. StringTie enables improved reconstruction of a transcriptome from RNA-seq reads. Nature Biotechnology 33(3): 290–5, Doi: 10.1038/nbt.3122.

37. Bolger, A.M., Lohse, M., Usadel, B., 2014. Trimmomatic : a flexible trimmer for Illumina sequence data. Bioinformatics 30(15): 2114–20, Doi: 10.1093/bioinformatics/btu170.

38. Salzberg, B.L. and S.L., 2013. Fast gapped-read alignment with Bowtie 2. Nature Methods 9(4): 357–9, Doi: 10.1038/nmeth.1923.Fast.

39. Ramírez, F., Ryan, D.P., Grüning, B., Bhardwaj, V., Kilpert, F., Richter, A.S., et al., 2016. deepTools2: a next generation web server for deep-sequencing data analysis. Nucleic Acids Research 44(W1): W160–5, Doi: 10.1093/nar/gkw257.

40. Liu, X., Wang, C., Liu, W., Li, J., Li, C., Kou, X., et al., 2016. Distinct features of H3K4me3 and H3K27me3 chromatin domains in pre-implantation embryos. Nature 537(7621): 558–62, Doi: 10.1038/nature19362.

41. Zhang, B., Zheng, H., Huang, B., Li, W., Xiang, Y., Peng, X., et al., 2016. Allelic reprogramming of the histone modification H3K4me3 in early mammalian development. Nature 537(7621): 553–7, Doi: 10.1038/nature19361.

42. Jung, Y.H., Sauria, M.E.G., Lyu, X., Cheema, M.S., Ausio, J., Taylor, J., et al., 2017. Chromatin States in Mouse Sperm Correlate with Embryonic and Adult Regulatory Landscapes. Cell Reports 18(6): 1366–82, Doi: 10.1016/j.celrep.2017.01.034.

43. Liu, L., Leng, L., Liu, C., Lu, C., Yuan, Y., Wu, L., et al., 2019. An integrated chromatin accessibility and transcriptome landscape of human pre-implantation embryos. Nature Communications 10(1): 364, Doi: 10.1038/s41467-018-08244-0.

44. Shen, Y., Yue, F., McCleary, D.F., Ye, Z., Edsall, L., Kuan, S., et al., 2012. A map of the cis-regulatory sequences in the mouse genome. Nature 488(7409): 116–20, Doi: 10.1038/nature11243.

45. Chu, A., Casero, D., Thamotharan, S., Wadehra, M., Cosi, A., Devaskar, S.U., 2019. The Placental Transcriptome in Late Gestational Hypoxia Resulting in Murine Intrauterine Growth Restriction Parallels Increased Risk of Adult Cardiometabolic Disease. Scientific Reports 9(1): 1243, Doi: 10.1038/s41598-018-37627-y.

46. Krueger, F., Andrews, S.R., 2016. SNPsplit: Allele-specific splitting of alignments between genomes with known SNP genotypes [version 1; referees: 3 approved]. F1000Research 5: 1–15, Doi: 10.12688/F1000RESEARCH.9037.1.

47. Waskom, M.L., 2021. Seaborn : statistical data visualization Statement of need. The Journal of Open Source Software 6: 1–4, Doi: 10.21105/joss.03021.

48. Harris, C.R., Millman, K.J., van der Walt, S.J., Gommers, R., Virtanen, P., Cournapeau, D., et al., 2020. Array programming with NumPy. Nature 585(7825): 357–62, Doi: 10.1038/s41586-020-2649-2.

49. Mckinney, W., 2010. Data Structures for Statistical Computing in Python. Proceedings of the 9th Python in Science Conference, vol. 1. p. 56–61.

50. Hunter, J.D., 2007. Matplotlib: A 2D Graphics Environment. Computing in Science & Engineering 9(3): 90–5, Doi: 10.1109/MCSE.2007.55.

51. Team, R.C., 2018. R: A language and environment for statistical computing.

52. Love, M.I., Huber, W., Anders, S., 2014. Moderated estimation of fold change and dispersion for RNA-seq data with DESeq2. Genome Biology 15(12): 1–21, Doi: 10.1186/s13059-014-0550-8.

53. Ignatiadis, N., Klaus, B., Zaugg, J.B., Huber, W., 2016. Data-driven hypothesis weighting increases detection power in genome-scale multiple testing. Nature Methods 13(7): 577–80, Doi: 10.1038/nmeth.3885.

54. Benjamini, Y., Hochberg, Y., 1995. Controlling the False Discovery Rate: A Practical and Powerful Approach to Multiple Testing. Journal of the Royal Statistical Society. Series B (Methodological) 57(1): 289–300.

55. Yi, L., Pimentel, H., Bray, N.L., Pachter, L., 2018. Gene-level differential analysis at transcript-level resolution. Genome Biology 19(1): 1–11, Doi: 10.1186/s13059-018-1419-z.

56. Ritchie, M.E., Phipson, B., Wu, D., Hu, Y., Law, C.W., Shi, W., et al., 2015. limma powers differential expression analyses for RNA-sequencing and microarray studies. Nucleic Acids Research 43(7): e47, Doi: 10.1093/nar/gkv007.

57. Taiyun, Wei, Simko, V., 2021. R package “corrplot”: Visualization of a Correlation Matrix.

58. Kolde, R., 2019. pheatmap: Pretty Heatmaps. R package.

59. Wickham, H., 2016. ggplot2: Elegant Graphics for Data Analysis. New York: Springer-Verlag.

60. Shen, L., 2014. GeneOverlap : An R package to test and visualize gene overlaps: 1–10.

61. Brionne, A., Juanchich, A., Hennequet-Antier, C., 2019. ViSEAGO: A Bioconductor package for clustering biological functions using Gene Ontology and semantic similarity. BioData Mining 12(1): 1–13, Doi: 10.1186/s13040-019-0204-1.

62. Lun, A.T.L., Smyth, G.K., 2016. csaw: a Bioconductor package for differential binding analysis of ChIP-seq data using sliding windows. Nucleic Acids Research 44(5): e45, Doi: 10.1093/nar/gkv1191.

63. Zhang, Y., Parmigiani, G., Johnson, W.E., 2020. ComBat-seq: batch effect adjustment for RNA-seq count data. NAR Genomics and Bioinformatics 2(3), Doi: 10.1093/nargab/lqaa078.

64. Leek, J.T., Johnson, W.E., Parker, H.S., Jaffe, A.E., Storey, J.D., 2012. The sva package for removing batch effects and other unwanted variation in high-throughput experiments. Bioinformatics (Oxford, England) 28(6): 882–3, Doi: 10.1093/bioinformatics/bts034.

65. Oksanen, J., Kindt, R., Legendre, P., O’Hara, B., Stevens, M.H.H., Oksanen, M.J., et al., 2007. The vegan package. Community Ecology Package 10(631–637): 719.

66. Welch, R.P., Lee, C., Imbriano, P.M., Patil, S., Weymouth, T.E., Smith, R.A., et al., 2014. ChIP-Enrich: gene set enrichment testing for ChIP-seq data. Nucleic Acids Research 42(13): e105– e105, Doi: 10.1093/nar/gku463.

67. Alexa, A., Rahnenführer, J., Lengauer, T., 2006. Improved scoring of functional groups from gene expression data by decorrelating GO graph structure. Bioinformatics 22(13): 1600–7, Doi: 10.1093/bioinformatics/btl140.

68. Cavalcante, R.G., Sartor, M.A., 2017. annotatr: genomic regions in context. Bioinformatics (Oxford, England) 33(15): 2381–3, Doi: 10.1093/bioinformatics/btx183.

69. Conway, J.R., Lex, A., Gehlenborg, N., 2017. UpSetR: an R package for the visualization of intersecting sets and their properties. Bioinformatics 33(18): 2938–40, Doi: 10.1093/bioinformatics/btx364.

70. Gel, B., Díez-Villanueva, A., Serra, E., Buschbeck, M., Peinado, M.A., Malinverni, R., 2016. regioneR: an R/Bioconductor package for the association analysis of genomic regions based on permutation tests. Bioinformatics 32(2): 289–91, Doi: 10.1093/bioinformatics/btv562.

71. Pohl, A., Beato, M., 2014. bwtool: a tool for bigWig files. Bioinformatics 30(11): 1618–9, Doi: 10.1093/bioinformatics/btu056.

72. Kim, S., Sohn, I., Ahn, J.-I., Lee, K.-H., Lee, Y.S., Lee, Y.S., 2004. Hepatic gene expression profiles in a long-term high-fat diet-induced obesity mouse model. Gene 340(1): 99–109, Doi: 10.1016/j.gene.2004.06.015.

73. Fei, X., Jiali, D., Yajie, G., Yuguo, N., Feixiang, Y., Junjie, Y., et al., 2016. BTG1 ameliorates liver steatosis by decreasing stearoyl-CoA desaturase 1 (SCD1) abundance and altering hepatic lipid metabolism. Science Signaling 9(428): ra50–ra50, Doi: 10.1126/scisignal.aad8581.

74. Støy, J., Kampmann, U., Mengel, A., Magnusson, N.E., Jessen, N., Grarup, N., et al., 2015. Reduced CD300LG mRNA tissue expression, increased intramyocellular lipid content and impaired glucose metabolism in healthy male carriers of Arg82Cys in CD300LG: a novel genometabolic cross-link between CD300LG and common metabolic phenotypes. BMJ Open Diabetes Research & Care 3(1): e000095, Doi: 10.1136/bmjdrc-2015-000095.

75. Perie, L., Verma, N., Mueller, E., 2021. The forkhead box transcription factor FoxP4 regulates thermogenic programs in adipocytes. Journal of Lipid Research 62, Doi: 10.1016/j.jlr.2021.100102.

76. Lacroix, M., Linares, L.K., Rueda-Rincon, N., Bloch, K., Di Michele, M., De Blasio, C., et al., 2021. The multifunctional protein E4F1 links P53 to lipid metabolism in adipocytes. Nature Communications 12(1): 7037, Doi: 10.1038/s41467-021-27307-3.

77. Lu, D., Xia, Q., Yang, Z., Gao, S., Sun, S., Luo, X., et al., 2021. ENO3 promoted the progression of NASH by negatively regulating ferroptosis via elevation of GPX4 expression and lipid accumulation. Annals of Translational Medicine; Vol 9, No 8 (April 2021): Annals of Translational Medicine.

78. Jia, X., Zhai, T., 2019. Integrated Analysis of Multiple Microarray Studies to Identify Novel Gene Signatures in Non-alcoholic Fatty Liver Disease. Frontiers in Endocrinology 10, Doi: 10.3389/fendo.2019.00599.

79. Chu, Y., Rosso, L.G., Huang, P., Wang, Z., Xu, Y., Yao, X., et al., 2014. Liver Med23 ablation improves glucose and lipid metabolism through modulating FOXO1 activity. Cell Research 24(10): 1250–65, Doi: 10.1038/cr.2014.120.

80. Iwasaki, H., 2009. Impaired PRMT1 activity in the liver and pancreas of type 2 diabetic Goto–Kakizaki rats. Life Sciences 85(3): 161–6, Doi: https://doi.org/10.1016/j.lfs.2009.05.007.

81. Choi, S., Choi, D., Lee, Y.-K., Ahn, S.H., Seong, J.K., Chi, S.W., et al., 2021. Depletion of Prmt1 in Adipocytes Impairs Glucose Homeostasis in Diet-Induced Obesity. Diabetes 70(8): 1664–78, Doi: 10.2337/db20-1050.

82. Pacana, T., Cazanave, S., Verdianelli, A., Patel, V., Min, H.-K., Mirshahi, F., et al., 2015. Dysregulated Hepatic Methionine Metabolism Drives Homocysteine Elevation in Diet-Induced Nonalcoholic Fatty Liver Disease. PloS One 10(8): e0136822, Doi: 10.1371/journal.pone.0136822.

83. Gerdes, P., Richardson, S.R., Mager, D.L., Faulkner, G.J., 2016. Transposable elements in the mammalian embryo: pioneers surviving through stealth and service. Genome Biology 17: 100, Doi: 10.1186/s13059-016-0965-5.

84. The Jackson Laboratory., 2022. Mouse Genome Informatics. http://www.informatics.jax.org.

85. Perez-Garcia, V., Fineberg, E., Wilson, R., Murray, A., Mazzeo, C.I., Tudor, C., et al., 2018. Placentation defects are highly prevalent in embryonic lethal mouse mutants. Nature 555(7697): 463–8, Doi: 10.1038/nature26002.

86. Wang, X., Miller, D.C., Harman, R., Antczak, D.F., Clark, A.G., 2013. Paternally expressed genes predominate in the placenta. Proceedings of the National Academy of Sciences 110(26): 10705–10, Doi: 10.1073/pnas.1308998110.

87. Chu, A., Casero, D., Thamotharan, S., Wadehra, M., Cosi, A., Devaskar, S.U., 2019. The Placental Transcriptome in Late Gestational Hypoxia Resulting in Murine Intrauterine Growth Restriction Parallels Increased Risk of Adult Cardiometabolic Disease. Scientific Reports 9(1): 1–15, Doi: 10.1038/s41598-018-37627-y.

88. Naruse, M., Ono, R., Irie, M., Nakamura, K., Furuse, T., Hino, T., et al., 2014. Sirh7/Ldoc1 knockout mice exhibit placental P4 overproduction and delayed parturition. Development (Cambridge, England) 141(24): 4763–71, Doi: 10.1242/dev.114520.

89. Henke, C., Strissel, P.L., Schubert, M.-T., Mitchell, M., Stolt, C.C., Faschingbauer, F., et al., 2015. Selective expression of sense and antisense transcripts of the sushi-ichi-related retrotransposon – derived family during mouse placentogenesis. Retrovirology 12(1): 9, Doi: 10.1186/s12977-015-0138-8.

90. Shan, N., Xiao, X., Chen, Y., Luo, X., Yin, N., Deng, Q., et al., 2016. Expression of DAB2IP in human trophoblast and its role in trophoblast invasion. The Journal of Maternal-Fetal & Neonatal Medicine : The Official Journal of the European Association of Perinatal Medicine, the Federation of Asia and Oceania Perinatal Societies, the International Society of Perinatal Obstetricians 29(3): 393–9, Doi: 10.3109/14767058.2014.1001974.

91. Perschbacher, K.J., Deng, G., Sandgren, J.A., Walsh, J.W., Witcher, P.C., Sapouckey, S.A., et al., 2020. Reduced mRNA Expression of RGS2 (Regulator of G Protein Signaling-2) in the Placenta Is Associated With Human Preeclampsia and Sufficient to Cause Features of the Disorder in Mice. Hypertension (Dallas, Tex. : 1979) 75(2): 569–79, Doi: 10.1161/HYPERTENSIONAHA.119.14056.

92. Yoshida, K., Maekawa, T., Ly, N.H., Fujita, S. ichiro., Muratani, M., Ando, M., et al., 2020. ATF7-Dependent Epigenetic Changes Are Required for the Intergenerational Effect of a Paternal Low-Protein Diet. Molecular Cell 78(3): 445–458.e6, Doi: 10.1016/j.molcel.2020.02.028.

93. Kimmins, S., Sassone-Corsi, P., 2005. Chromatin remodelling and epigenetic features of germ cells. Nature 434(7033): 583–9, Doi: 10.1038/nature03368.

94. Larose, H., Shami, A.N., Abbott, H., Manske, G., Lei, L., Hammoud, S.S., 2019. Gametogenesis: A journey from inception to conception. Current Topics in Developmental Biology 132: 257–310, Doi: 10.1016/bs.ctdb.2018.12.006.

95. Lambrot, R., Lafleur, C., Kimmins, S., 2015. The histone demethylase KDM1A is essential for the maintenance and differentiation of spermatogonial stem cells and progenitors. FASEB Journal 29(11): 4402–16, Doi: 10.1096/fj.14-267328.

96. Maezawa, S., Yukawa, M., Alavattam, K.G., Barski, A., Namekawa, S.H., 2018. Dynamic reorganization of open chromatin underlies diverse transcriptomes during spermatogenesis. Nucleic Acids Research 46(2): 593–608, Doi: 10.1093/nar/gkx1052.

97. Erkek, S., Hisano, M., Liang, C.Y., Gill, M., Murr, R., Dieker, J., et al., 2013. Molecular determinants of nucleosome retention at CpG-rich sequences in mouse spermatozoa. Nature Structural and Molecular Biology 20(7): 868–75, Doi: 10.1038/nsmb.2599.

98. Hammoud, S.S., Nix, D.A., Zhang, H., Purwar, J., Carrell, D.T., Cairns, B.R., 2009. Distinctive chromatin in human sperm packages genes for embryo development. Nature 460(7254): 473–8, Doi: 10.1038/nature08162.

99. Lesch, B.J., Silber, S.J., McCarrey, J.R., Page, D.C., 2016. Parallel evolution of male germline epigenetic poising and somatic development in animals. Nature Genetics 48(8): 888–94, Doi: 10.1038/ng.3591.

100. Brykczynska, U., Hisano, M., Erkek, S., Ramos, L., Oakeley, E.J., Roloff, T.C., et al., 2010. Repressive and active histone methylation mark distinct promoters in human and mouse spermatozoa. Nature Structural and Molecular Biology 17(6): 679–87, Doi: 10.1038/nsmb.1821.

101. Lambrot, R., Chan, D., Shao, X., Aarabi, M., Kwan, T., Bourque, G., et al., 2021. Whole-genome sequencing of H3K4me3 and DNA methylation in human sperm reveals regions of overlap linked to fertility and development. Cell Reports 36(3): 109418, Doi: https://doi.org/10.1016/j.celrep.2021.109418.

102. Sánchez-Margalet, V., Valle, M., Ruz, F.J., Gascón, F., Mateo, J., Goberna, R., 2002. Elevated plasma total homocysteine levels in hyperinsulinemic obese subjects. The Journal of Nutritional Biochemistry 13(2): 75–9, Doi: https://doi.org/10.1016/S0955-2863(01)00197-8.

103. Karatela, R.A., Sainani, G.S., 2009. Plasma homocysteine in obese, overweight and normal weight hypertensives and normotensives. Indian Heart Journal 61(2): 156–9.

104. Morgan, H.D., Sutherland, H.G.E., Martin, D.I.K., Whitelaw, E., 1999. Epigenetic inheritance at the agouti locus in the mouse. Nature Genetics 23(3): 314–8, Doi: 10.1038/15490.

105. Dolinoy, D.C., Weidman, J.R., Waterland, R.A., Jirtle, R.L., 2006. Maternal genistein alters coat color and protects Avy mouse offspring from obesity by modifying the fetal epigenome. Environmental Health Perspectives 114(4): 567–72, Doi: 10.1289/ehp.8700.

106. Rakyan, V.K., Chong, S., Champ, M.E., Cuthbert, P.C., Morgan, H.D., Luu, K.V.K., et al., 2003. Transgenerational inheritance of epigenetic states at the murine AxinFu allele occurs after maternal and paternal transmission. Proceedings of the National Academy of Sciences of the United States of America 100(5): 2538–43, Doi: 10.1073/pnas.0436776100.

107. Padmanabhan, N., Jia, D., Geary-joo, C., Wu, X., Ferguson-smith, A.C., Fung, E., et al., 2013. Mutation in Folate Metabolism Causes Epigenetic Instability and Transgenerational Effects on Development. Cell 155(1): 81–93, Doi: 10.1016/j.cell.2013.09.002.

108. Anway, M.D., Cupp, A.S., Uzumcu, M., Skinner, M.K., 2005. Epigenetic Transgenerational Actions of Endocrine Disruptors and Male Fertility. Science 308(5727): 1466 LP – 1469, Doi: 10.1126/science.1108190.

109. Martínez, D., Pentinat, T., Ribó, S., Daviaud, C., Bloks, V.W., Cebrià, J., et al., 2014. In utero undernutrition in male mice programs liver lipid metabolism in the second-generation offspring involving altered Lxra DNA methylation. Cell Metabolism 19(6): 941–51, Doi: 10.1016/j.cmet.2014.03.026.

110. Wei, Y., Yang, C.-R., Wei, Y.-P., Zhao, Z.-A., Hou, Y., Schatten, H., et al., 2014. Paternally induced transgenerational inheritance of susceptibility to diabetes in mammals. Proceedings of the National Academy of Sciences 111(5): 1873–8, Doi: 10.1073/pnas.1321195111.

111. Carone, B.R., Fauquier, L., Habib, N., Shea, J.M., Caroline, E., Li, R., et al., 2011. Paternally-induced transgenerational environmental reprogramming of metabolic gene expression in mammals Benjamin 143(7): 1084–96, Doi: 10.1016/j.cell.2010.12.008.Paternally-induced.

112. Kerr, B., Leiva, A., Farías, M., Contreras-Duarte, S., Toledo, F., Stolzenbach, F., et al., 2018. Foetoplacental epigenetic changes associated with maternal metabolic dysfunction. Placenta 69: 146–52, Doi: 10.1016/j.placenta.2018.04.006.

113. Franzago, M., Fraticelli, F., Stuppia, L., Vitacolonna, E., 2019. Nutrigenetics , epigenetics and gestational diabetes : consequences in mother and child. Epigenetics 14(3): 215–35, Doi: 10.1080/15592294.2019.1582277.

114. Moore, T., 2001. Genetic conflict, genomic imprinting and establishment of the epigenotype in relation to growth. Reproduction (Cambridge, England) 122(2): 185–93, Doi: 10.1530/rep.0.1220185.

115. Binder, N.K., Hannan, N.J., Gardner, D.K., 2012. Paternal Diet-Induced Obesity Retards Early Mouse Embryo Development, Mitochondrial Activity and Pregnancy Health. PLoS ONE 7(12), Doi: 10.1371/journal.pone.0052304.

116. Rosenfeld, C.S., 2015. Sex-Specific Placental Responses in Fetal Development. Endocrinology 156(10): 3422–34, Doi: 10.1210/en.2015-1227.

117. Naruse, K., Tsunemi, T., Kawahara, N., Kobayashi, H., 2019. Preliminary evidence of a paternal-maternal genetic conflict on the placenta: Link between imprinting disorder and multi-generational hypertensive disorders. Placenta 84: 69–73, Doi: https://doi.org/10.1016/j.placenta.2019.02.009.

118. Denomme, M.M., Haywood, M.E., Parks, J.C., Schoolcraft, W.B., Katz-Jaffe, M.G., 2020. The inherited methylome landscape is directly altered with paternal aging and associated with offspring neurodevelopmental disorders. Aging Cell 19(8): e13178, Doi: https://doi.org/10.1111/acel.13178.

119. Denomme, M.M., Parks, J.C., McCallie, B.R., McCubbin, N.I., Schoolcraft, W.B., Katz-Jaffe, M.G., 2020. Advanced paternal age directly impacts mouse embryonic placental imprinting. PLoS ONE 15(3): 1–13, Doi: 10.1371/journal.pone.0229904.

120. Gabory, A., Attig, L., Junien, C., 2011. Developmental programming and epigenetics. The American Journal of Clinical Nutrition 94(suppl_6): 1943S–1952S, Doi: 10.3945/ajcn.110.000927.

121. Nelissen, E.C.M., van Montfoort, A.P.A., Dumoulin, J.C.M., Evers, J.L.H., 2011. Epigenetics and the placenta. Human Reproduction Update 17(3): 397–417, Doi: 10.1093/humupd/dmq052.

122. Binder, N.K., Beard, S.A., Kaitu’U-Lino, T.J., Tong, S., Hannan, N.J., Gardner, D.K., 2015. Paternal obesity in a rodent model affects placental gene expression in a sex-specific manner. Reproduction 149(5): 435–44, Doi: 10.1530/REP-14-0676.

123. Jazwiec, P.A., Patterson, V.S., Ribeiro, T.A., Yeo, E., Kennedy, K.M., Mathias, P.C.F., et al., 2021. Paternal obesity results in placental hypoxia and sex-specific impairments in placental vascularization and offspring metabolic function. BioRxiv: 2021.03.27.437284, Doi: 10.1101/2021.03.27.437284.

124. Leavey, K., Wilson, S.L., Bainbridge, S.A., Robinson, W.P., Cox, B.J., 2018. Epigenetic regulation of placental gene expression in transcriptional subtypes of preeclampsia. Clinical Epigenetics 10(1): 28, Doi: 10.1186/s13148-018-0463-6.

125. Sun, N., Chen, H., Ma, Y., Pang, W., Wang, X., Zhang, Q., 2020. H3K4me3-Mediated Upregulation of LncRNA-HEIPP in Preeclampsia Placenta Affects Invasion of Trophoblast Cells 11(December): 1–11, Doi: 10.3389/fgene.2020.559478.

126. Meister, S., Hahn, L., Beyer, S., Kuhn, C., Jegen, M., von Schönfeldt, V., et al., 2021. Epigenetic modification via H3K4me3 and H3K9ac in human placenta is reduced in preeclampsia. Journal of Reproductive Immunology 145: 103287, Doi: https://doi.org/10.1016/j.jri.2021.103287.

127. Eriksson, J.G., Kajantie, E., Osmond, C., Thornburg, K., Barker, D.J.P., 2010. Boys live dangerously in the womb. American Journal of Human Biology : The Official Journal of the Human Biology Council 22(3): 330–5, Doi: 10.1002/ajhb.20995.

128. Gupte, A.A., Pownall, H.J., Hamilton, D.J., 2015. Estrogen : An Emerging Regulator of Insulin Action and Mitochondrial Function 2015.

129. Barratt, C.L.R., De Jonge, C.J., Anderson, R.A., Eisenberg, M.L., Garrido, N., Rautakallio Hokkanen, S., et al., 2021. A global approach to addressing the policy, research and social challenges of male reproductive health. Human Reproduction Open 2021(1): hoab009, Doi: 10.1093/hropen/hoab009.

