## Supplementary material for "Sperm Histone H3 Lysine 4 tri-methylation serves as a metabolic sensor of paternal obesity and is associated with the inheritance of metabolic dysfunction": Tables S1-4 and S8

**Supplemental tables**

**Table S1. Diets' energy density and macronutrients composition**

| | **Control diet** (CON; D12450J, Research Diets Inc.) | **High-fat diet** (HFD; D12492, Research Diets Inc.) | **Regular chow diet** (2020X, Teklad Diets) |
|---|---|---|---|
| **Energy density (kcal/g)** | 3.85 | 5.24 | 3.1 |
| **Calories from Protein (%)** | 20 | 20 | 24 |
| **Calories from Fat (%)** | 10 | 60 | 16 |
| **Calories from Carbohydrate (%)** | 70 | 20 | 60 |

**Table S2. Number of animals used per group per sex per generation for metabolic characterization**

| | **$F_0$ males** | **$F_1$ males** | **$F_2$ males** | **$F_1$ females** | **$F_2$ females** |
|---|---|---|---|---|---|
| **WT CON** | 17 | 35 | 18 | 38 | 15 |
| **WT HFD** | 18 | 28 | 19 | 39 | 19 |
| **TG CON** | 15 | 30 | 8 | 49 | 13 |
| **TG HFD** | 25 | 43 | 11 | 42 | 21 |

**Table S3. Litter size generated by $F_0$ and $F_1$ sires**

| **Group** | **WT CON** | **TG CON** | **WT HFD** | **TG HFD** | **Significance** |
|---|---|---|---|---|---|
| **$F_0$ litter size Mean ± SEM (N=)** | 6.833 ± 0.458 (12) | 7.9 ± 0.767 (10) | 6.333 ± 0.607 (12) | 6.2 ± 0.48 (15) | NS |
| **$F_1$ litter size Mean ± SEM (N=)** | 7 ± 0.5 (8) | 4.6 ± 1.288 (5) | 5.667 ± 0.689 (12) | 4.875 ± 0.666 (8) | NS |

**Table S4. Sex ratios of litters generated by $F_0$ and $F_1$ sires**

| **Group** | **WT CON** | **TG CON** | **WT HFD** | **TG HFD** | **Significance** |
|---|---|---|---|---|---|
| **$F_0$ sex ratio Mean ± SEM (N=)** | 0.509 ± 0.068 (12) | 0.343 ± 0.069 (10) | 0.534 ± 0.071 (12) | 0.579 ± 0.055 (15) | NS |
| **$F_1$ sex ratio Mean ± SEM (N=)** | 0.476 ± 0.072 (8) | 0.548 ± 0.141 (5) | 0.598 ± 0.073 (12) | 0.425 ± 0.095 (8) | NS |

**Table S8. Sperm H3K4me3 ChIP-Sequencing read numbers and alignment rates**

| Batch | Diet | Genotype | Reads # | Alignment rate (%) |
|---|---|---|---|---|
| 1 | CON | WT | 25,237,875 | 91.24 |
| 2 | CON | WT | 36,679,172 | 98.04 |
| 3 | CON | WT | 44,156,311 | 97.58 |
| 4 | CON | WT | 38,188,136 | 97.57 |
| 5 | CON | WT | 32,891,823 | 97.25 |
| 1 | HFD | WT | 30,356,006 | 94.03 |
| 2 | HFD | WT | 36,934,484 | 97.11 |
| 3 | HFD | WT | 31,718,194 | 97.38 |
| 4 | HFD | WT | 27,841,763 | 97.9 |
| 5 | HFD | WT | 30,902,900 | 96.91 |
| 1 | CON | TG | 29,310,441 | 96.88 |
| 2 | CON | TG | 38,274,515 | 96.88 |
| 3 | CON | TG | 27,291,467 | 97.27 |
| 4 | CON | TG | 39,918,514 | 97.81 |
| 5 | CON | TG | 30,925,671 | 97.68 |
| 1 | HFD | TG | 23,856,350 | 94.75 |
| 2 | HFD | TG | 39,534,589 | 97.93 |
| 3 | HFD | TG | 38,469,968 | 97.73 |
| 4 | HFD | TG | 28,052,094 | 98.14 |
| 5 | HFD | TG | 35,028,161 | 98.13 |
|  |  | **Average** | 33,278,421.7 | 96.9105 |
