## Supplementary material for "Sperm Histone H3 Lysine 4 tri-methylation serves as a metabolic sensor of paternal obesity and is associated with the inheritance of metabolic dysfunction": Figures S1-7

**Figure S1**

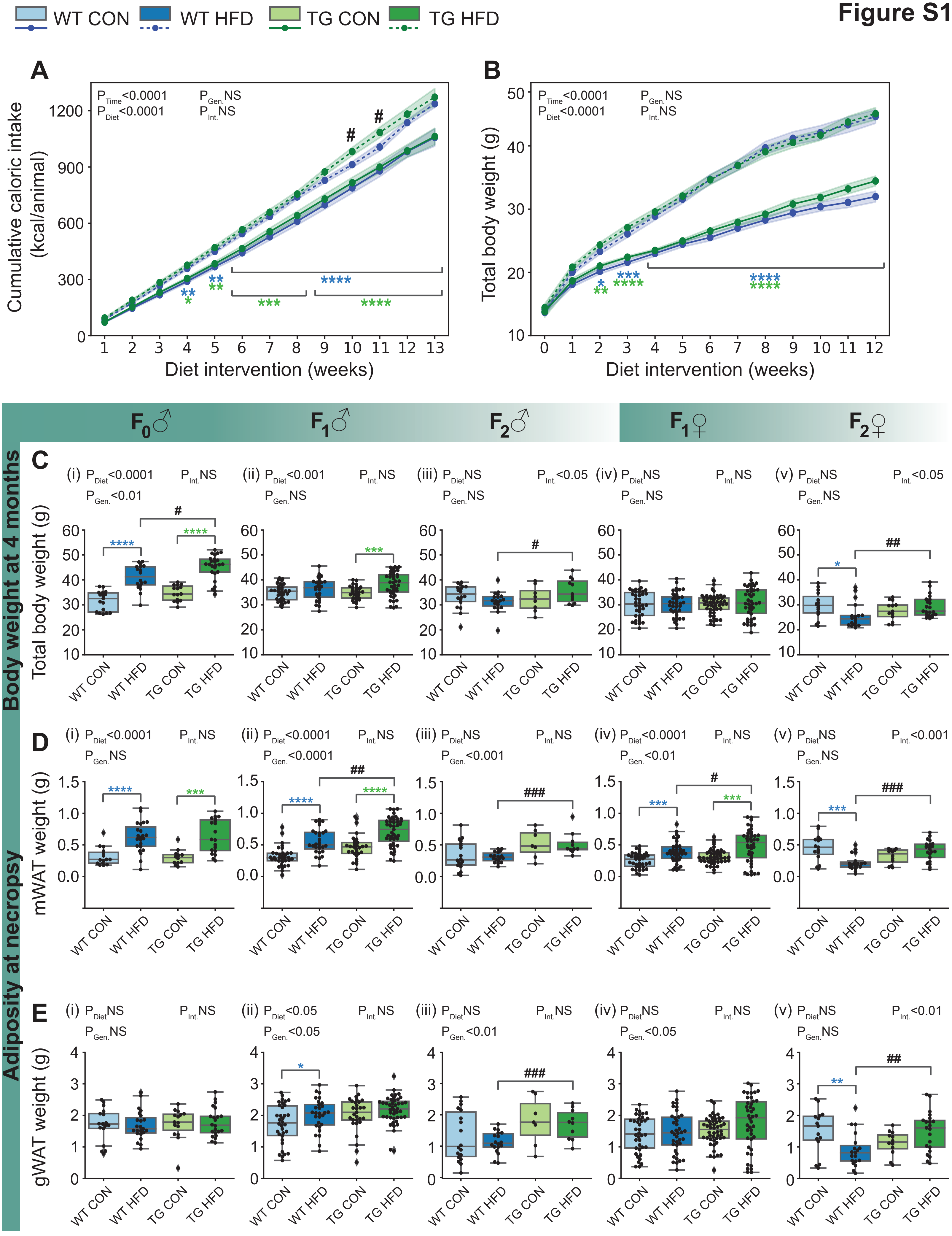
WT CON
WT HFD
TG CON
TG HFD
A
P Time <0.0001
P Diet <0.0001
P Gen. NS
P Int. NS
Cumulative caloric intake (kcal/animal)
Diet intervention (weeks)
B
P Time <0.0001
P Diet <0.0001
P Gen. NS
P Int. NS
Total body weight (g)
Diet intervention (weeks)
F0♂
F1♂
F2♂
F1♀
F2♀
Body weight at 4 months
C
(i) P Diet <0.0001 P Int. NS P Gen. <0.01
(ii) P Diet <0.001 P Int. NS P Gen. NS
(iii) P Diet NS P Int. <0.05 P Gen. NS
(iv) P Diet NS P Int. NS P Gen. NS
(v) P Diet NS P Int. <0.05 P Gen. NS
Total body weight (g)
Adiposity at necropsy
D
(i) P Diet <0.0001 P Int. NS P Gen. NS
(ii) P Diet <0.0001 P Int. NS P Gen. <0.0001
(iii) P Diet NS P Int. NS P Gen. <0.001
(iv) P Diet <0.0001 P Int. NS P Gen. <0.01
(v) P Diet NS P Int. <0.001 P Gen. NS
mWAT weight (g)
E
(i) P Diet NS P Int. NS P Gen. NS
(ii) P Diet <0.05 P Int. NS P Gen. <0.05
(iii) P Diet NS P Int. NS P Gen. <0.01
(iv) P Diet NS P Int. NS P Gen. <0.05
(v) P Diet NS P Int. <0.01 P Gen. NS
gWAT weight (g)
WT CON
WT HFD
TG CON
TG HFD

**Figure S1: Paternal obesity increases body weight and fat accruement across generations** A) Cumulative energy intake during the dietary treatment. The amount of food consumed weekly per cage was measured and the cumulative caloric intake per mouse was calculated based on the calorie content specific to each diet. B) Growth trajectories of $F_0$ sires before and during the 12-week diet intervention. C) Total body weight at 4 months of age in $F_0$ males (i), $F_1$ males (ii), $F_2$ males (iii), $F_1$ females (iv) and $F_2$ females (v). D) Mesenteric white adipose tissue (mWAT) weight at necropsy in $F_0$ males (i), $F_1$ males (ii), $F_2$ males (iii), $F_1$ females (iv) and $F_2$ females (v). E) Gonadal white adipose tissue (gWAT) weight at necropsy in $F_0$ males (i), $F_1$ males (ii), $F_2$ males (iii), $F_1$females (iv) and $F_2$ females (v). Results are shown as mean ± SEM. Significance for main effects of diet, genotype, time, and for diet-genotype interactions are shown above each graph. Significance for pairwise comparisons are shown as the following: *P<0.05, **P<0.01, ***P<0.001, ****P<0.0001 (in blue; WT CON vs WT HFD, in green; TG CON vs TG HFD) and $^{\#}$P<0.05, $^{\#\#}$P<0.01, $^{\#\#\#}$P<0.001 (WT HFD vs TG HFD).

**Figure S2**

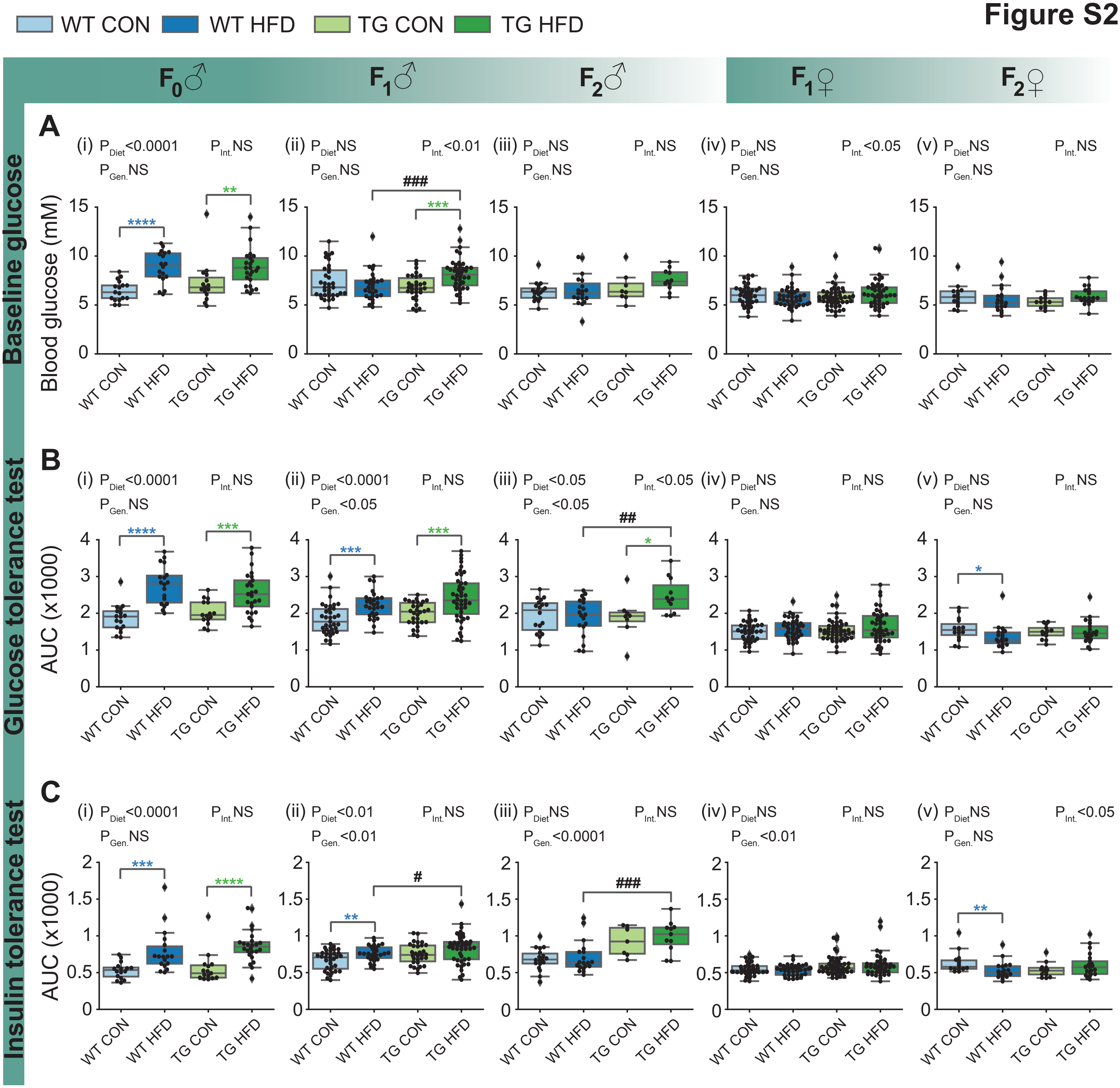
WT CON
WT HFD
TG CON
TG HFD
F0♂
F1♂
F2♂
F1♀
F2♀
A
Baseline glucose
Blood glucose (mM)
(i) P Diet <0.0001 P Int. NS P Gen. NS
(ii) P Diet NS P Int. <0.01 P Gen. NS
(iii) P Diet NS P Int. NS P Gen. NS
(iv) P Diet NS P Int. <0.05 P Gen. NS
(v) P Diet NS P Int. NS P Gen. NS
B
Glucose tolerance test
AUC (x1000)
(i) P Diet <0.0001 P Int. NS P Gen. NS
(ii) P Diet <0.0001 P Int. NS P Gen. <0.05
(iii) P Diet <0.05 P Int. <0.05 P Gen. <0.05
(iv) P Diet NS P Int. NS P Gen. NS
(v) P Diet NS P Int. NS P Gen. NS
C
Insulin tolerance test
AUC (x1000)
(i) P Diet <0.0001 P Int. NS P Gen. NS
(ii) P Diet <0.01 P Int. NS P Gen. <0.01
(iii) P Diet NS P Int. NS P Gen. <0.0001
(iv) P Diet NS P Int. NS P Gen. <0.01
(v) P Diet NS P Int. <0.05 P Gen. NS

**Figure S2: Paternal obesity alters metabolic profiles across generations in a sex-specific manner** A) Baseline blood glucose levels after overnight fasting (15 ±1 hour) at 4 months of age in $F_0$ males (i), $F_1$males (ii), $F_2$males (iii), $F_1$ females (iv) and $F_2$ males (v). B) Glucose tolerance test area under the curve (AUC) for $F_0$ males (i), $F_1$ males (ii), $F_2$ males (iii), $F_1$ females (iv) and $F_2$ females (v). C) Insulin tolerance test AUC for $F_0$ males (i), $F_1$ males (ii), $F_2$ males (iii), $F_1$females (iv) and $F_2$ females (v). The AUC was calculated using the trapezoidal rule from individual animal glucose tolerance test curves (in Fig. 1D) and insulin tolerance test curves (in Fig. 1E). Results are shown as mean ± SEM. Significance for main effects of diet, genotype, and for diet-genotype interactions are shown above each graph. Significance for pairwise comparisons are shown as the following: *P<0.05, **P<0.01, ***P<0.001, ****P<0.0001 (in blue; WT CON vs WT HFD, in green; TG CON vs TG HFD) and $^{\#}$P<0.05, $^{\#\#}$P<0.01 (WT HFD vs TG HFD).

**Figure S3**

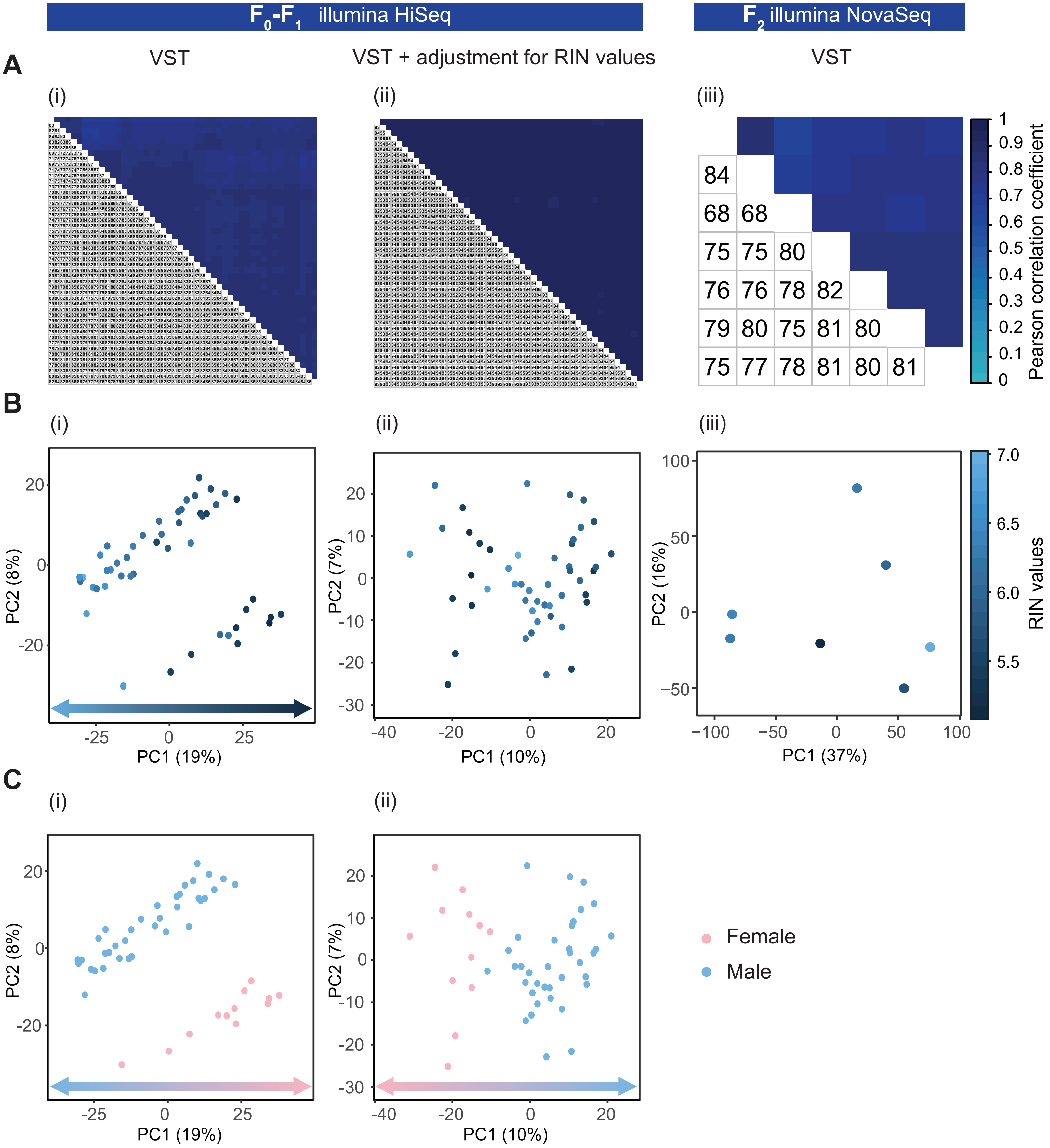

F0-F1 illumina HiSeq
F2 illumina NovaSeq
A
VST
VST + adjustment for RIN values
VST
(i)
(ii)
(iii)
84
68 68
75 75 80
76 76 78 82
79 80 75 81 80
75 77 78 81 80 81
Pearson correlation coefficient
1
0.9
0.8
0.7
0.6
0.5
0.4
0.3
0.2
0.1
0
B
(i)
(ii)
(iii)
PC2 (8%)
PC1 (19%)
PC2 (7%)
PC1 (10%)
PC2 (16%)
PC1 (37%)
RIN values
7.0
6.5
6.0
5.5
C
(i)
(ii)
PC2 (8%)
PC1 (19%)
PC2 (7%)
PC1 (10%)
Female
Male

**Figure S3: Liver RNA-sequencing data quality assessment and normalization** A) Pearson correlation heatmaps on transcripts with variance stabilizing transformation (VST), before (i) and after (ii) correcting for RIN values in $F_0$ and $F_1$ samples run on an illumina HiSeq platform, and in $F_2$ samples run on an illumina NovaSeq platform (iii). Color gradients indicate the Pearson correlation coefficients for each pairwise comparison of samples. B) Principal component analysis on transcripts with variance stabilizing transformation (VST), with samples labeled by RIN value before (i) and after (ii) correcting for RINs in $F_0$ and $F_1$ samples (illumina HiSeq) and in F2 samples (illumina NovaSeq) (iii). C) Principal component analysis on transcripts with variance stabilizing transformation (VST), with samples labeled by sex before (i) and after (ii) correcting for RINs in $F_0$ and $F_1$ samples (illumina HiSeq).

**Figure S4**

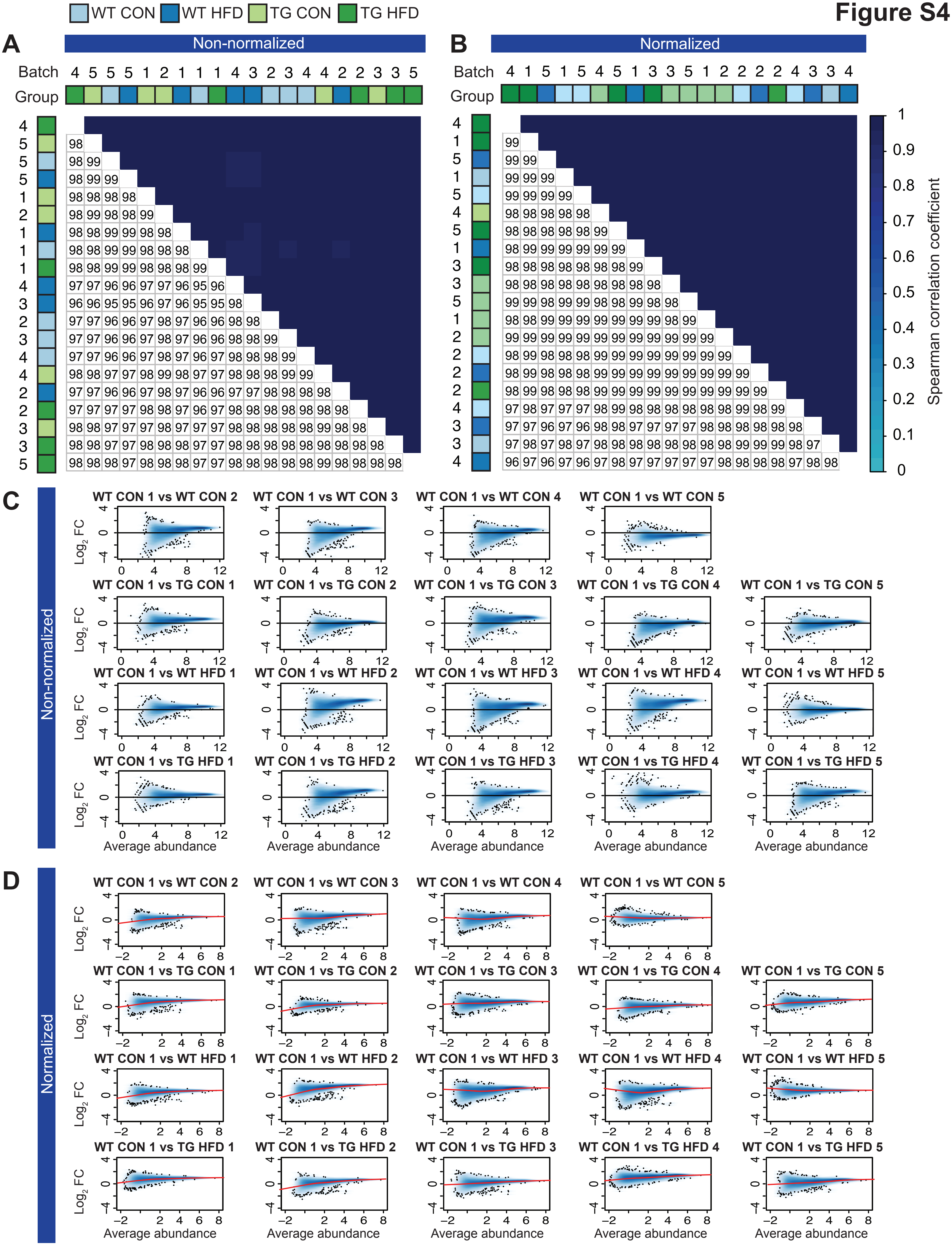
WT CON
WT HFD
TG CON
TG HFD
A
Non-normalized
B
Normalized
Batch
Group
Spearman correlation coefficient
C
Non-normalized
WT CON 1 vs WT CON 2
WT CON 1 vs WT CON 3
WT CON 1 vs WT CON 4
WT CON 1 vs WT CON 5
WT CON 1 vs TG CON 1
WT CON 1 vs TG CON 2
WT CON 1 vs TG CON 3
WT CON 1 vs TG CON 4
WT CON 1 vs TG CON 5
WT CON 1 vs WT HFD 1
WT CON 1 vs WT HFD 2
WT CON 1 vs WT HFD 3
WT CON 1 vs WT HFD 4
WT CON 1 vs WT HFD 5
WT CON 1 vs TG HFD 1
WT CON 1 vs TG HFD 2
WT CON 1 vs TG HFD 3
WT CON 1 vs TG HFD 4
WT CON 1 vs TG HFD 5
Log2 FC
Average abundance
D
Normalized
WT CON 1 vs WT CON 2
WT CON 1 vs WT CON 3
WT CON 1 vs WT CON 4
WT CON 1 vs WT CON 5
WT CON 1 vs TG CON 1
WT CON 1 vs TG CON 2
WT CON 1 vs TG CON 3
WT CON 1 vs TG CON 4
WT CON 1 vs TG CON 5
WT CON 1 vs WT HFD 1
WT CON 1 vs WT HFD 2
WT CON 1 vs WT HFD 3
WT CON 1 vs WT HFD 4
WT CON 1 vs WT HFD 5
WT CON 1 vs TG HFD 1
WT CON 1 vs TG HFD 2
WT CON 1 vs TG HFD 3
WT CON 1 vs TG HFD 4
WT CON 1 vs TG HFD 5
Log2 FC
Average abundance

**Figure S4: Sperm ChIP-sequencing data quality assessment and normalization** A-B) Spearman correlation heatmaps for genomic regions enriched with H3K4me3, before (A) and after (B) TMM normalization and batch correction. Colored boxes indicate sample groups (light blue=WT CON, dark blue=WT HFD, light green=TG CON, dark green=TG HFD) and numbers (from 1 to 5) indicate the sample batch. Color gradients indicate the Spearman correlation coefficients for each pairwise comparison of samples. C-D) MA-plots of pairwise comparisons between WT CON (rep 1) and all other samples, before (C) and after (D) TMM normalization and batch correction.

Figure S5

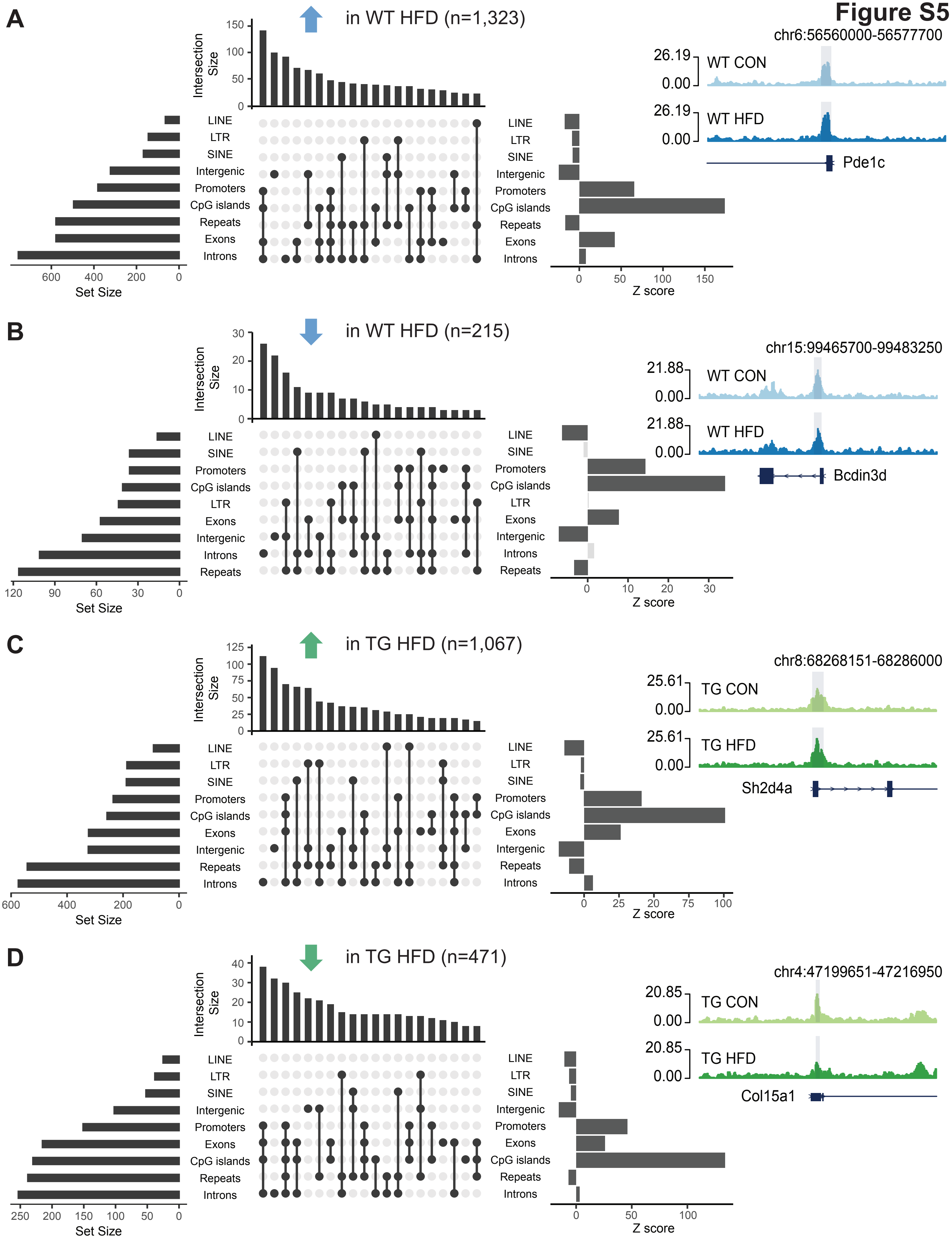
A
in WT HFD (n=1,323)
Intersection Size
Set Size
Z score
LINE
LTR
SINE
Intergenic
Promoters
CpG islands
Repeats
Exons
Introns
chr6:56560000-56577700
WT CON
WT HFD
Pde1c
B
in WT HFD (n=215)
chr15:99465700-99483250
Bcdin3d
C
in TG HFD (n=1,067)
chr8:68268151-68286000
TG CON
TG HFD
Sh2d4a
D
in TG HFD (n=471)
chr4:47199651-47216950
Col15a1

**Figure S5: Obesity-sensitive H3K4me3 regions are predominantly located in CpG islands, promoters, exons, and intergenic regions** A-D) Upset plots show genome annotation identifying the functional regions with obesity-induced differential enrichment of H3K4me3 in sperm according to directionality change, with increased enrichment in WT HFD (A), decreased enrichment in TG HFD (B), increased enrichment in TG HFD (C) and decreased enrichment in TG HFD (D). Horizontal bars on the left represent the number of regions belonging to each genomic annotation (set size). Vertical bars represent the number of regions belonging to intersecting annotations (intersection size). Intersection sets are represented by connecting nodes. Horizontal bars on the right represent the enrichment (z-score) for each respective annotation compared to what would be expected by chance if regions of similar sizes were randomly located across the genome ($p<0.05$, 1000 permutations). Dark grey bars represent significant enrichment whereas light grey bars are not significant. Genome browser snapshots show genes with deH3K4me3 in sperm (WT CON light blue, WT HFD dark blue, TG CON light green and TG HFD dark green).

### Figure S6

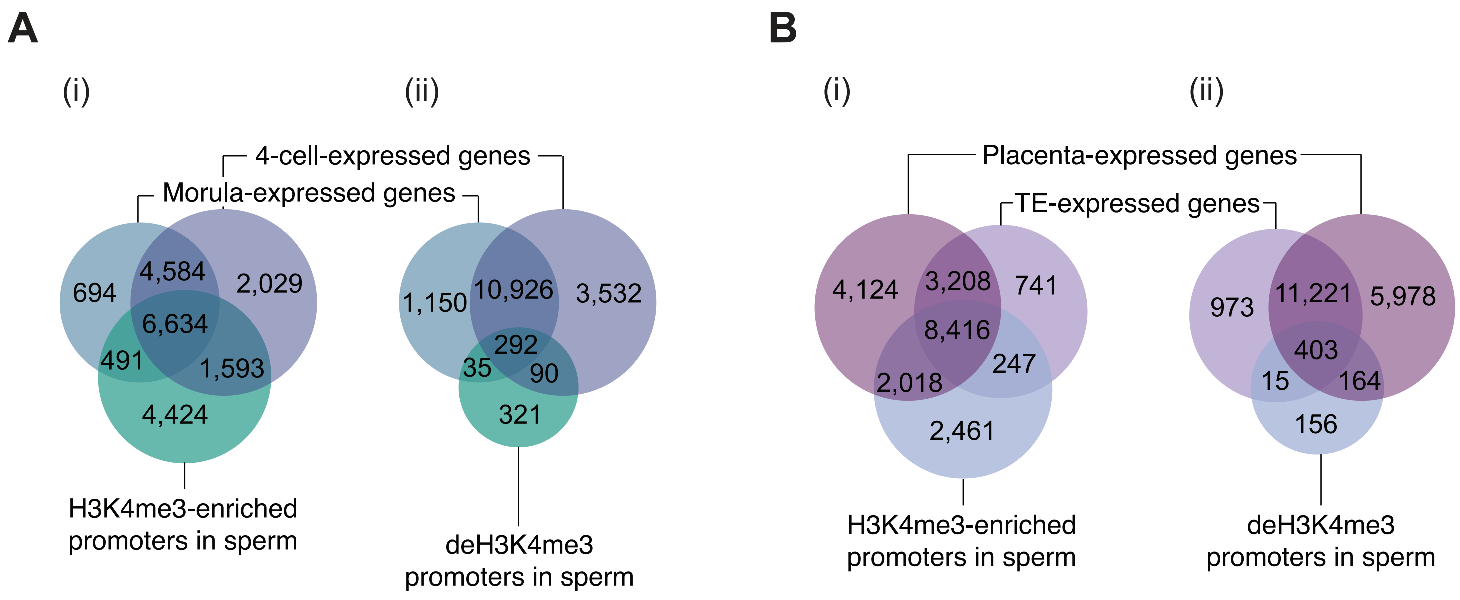
A
(i)
(ii)
4-cell-expressed genes
Morula-expressed genes
694
4,584
2,029
6,634
491
1,593
4,424
H3K4me3-enriched promoters in sperm
1,150
10,926
3,532
292
35
90
321
deH3K4me3 promoters in sperm
B
(i)
(ii)
Placenta-expressed genes
TE-expressed genes
4,124
3,208
741
8,416
247
2,018
2,461
H3K4me3-enriched promoters in sperm
973
11,221
5,978
403
15
164
156
deH3K4me3 promoters in sperm

**Figure S6: Obesity alters sperm H3K4me3 at genes expressed in the 4-cell and morula embryos, trophectoderm and placenta** A) Venn diagrams showing the overlap between genes expressed in the 4-cell embryo and genes expressed in the morula embryo, with genes with H3K4me3-enriched promoters in sperm (i) or with genes with diet-induced deH3K4me3 at promoters in sperm (ii). B) Venn diagrams showing the overlap between genes expressed in the trophectoderm and genes expressed in the placenta, with genes with H3K4me3-enriched promoters in sperm (i) or with genes with diet-induced deH3K4me3 at promoters in sperm (ii).

### Figure S7

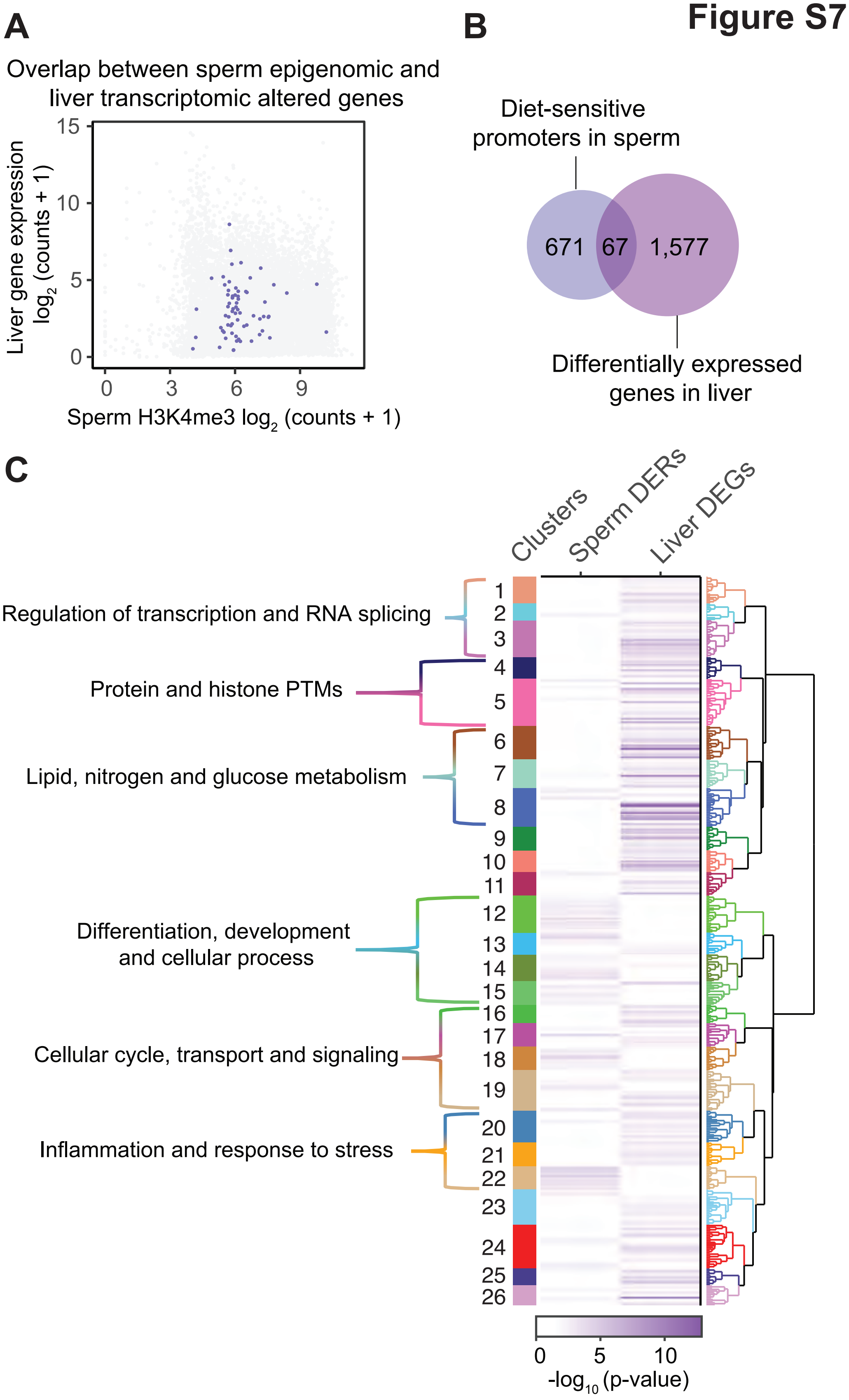
A
Overlap between sperm epigenomic and liver transcriptomic altered genes
Liver gene expression log2 (counts + 1)
Sperm H3K4me3 log2 (counts + 1)
0
3
6
9
5
10
15
B
Diet-sensitive promoters in sperm
671
67
1,577
Differentially expressed genes in liver
C
Clusters
Sperm DERs
Liver DEGs
Regulation of transcription and RNA splicing
Protein and histone PTMs
Lipid, nitrogen and glucose metabolism
Differentiation, development and cellular process
Cellular cycle, transport and signaling
Inflammation and response to stress
-log10 (p-value)

**Figure S7: Obesity-induced changes in H3K4me3 enrichment in sperm show minor overlap with genes altered in adult offspring liver** A) Scatterplot showing liver RNA expression values (y axis; log2 counts +1) and sperm H3K4me3 enrichment values (x axis; log2 counts + 1) for genes with paternal-diet induced differential expression in livers of $F_1$ males overlapping with deH3K4me3 at promoters in sperm. B) Venn diagram showing the overlap of genes enriched with diet-induced deH3K4me3 at promoters in sperm and genes with paternal-diet induced differential expression in livers of $F_1$ males. C) Heatmap of significant GO terms, comparing enriched biological functions in diet-induced sperm differentially enriched regions at promoters and liver differentially expressed genes in $F_1$ males WT and TG HFD. Rows represent enriched GO terms which are ordered by hierarchical clustering based on Wang's semantic similarity distance and *ward.D2* aggregation criterion. Each column represents a comparison of interest for which enriched GO terms were annotated based on the list of significant genes. The color gradient depicts the GO term enrichment significance ($-\log_{10}$ p-value). An interactive version of this heatmap can be found in Supplemental file 5 and the complete list of significantly enriched GO terms can be found in Table S22.
